# Aging: an inevitable road toward gut microbiota pathoadaptation

**DOI:** 10.1101/2025.01.29.635424

**Authors:** Rita Melo-Miranda, Ana Pinto, Hugo C. Barreto, Catarina Jesus, Isabel Gordo, Iola Duarte, Ana Sousa

## Abstract

Laboratory-raised mice live approximately seven times longer and healthier lives compared to their wild counterparts, due to a standardized healthy diet and limited exposure to environmental stressors^1^.

Aging is associated with increased inflammation and microbial dysbiosis^2–4^. Collectively, these influence microbiota evolution and may contribute to the enrichment in pathobiont frequency observed in old age^4^. Alternatively, this increase could stem from a decline in colonization resistance^5,6^, creating favorable conditions for pathobiont invasion.

Here, we sought to test whether aging in healthy, controlled conditions, could prevent the selection of age-associated pathobionts.

We have followed the adaptive evolution of a commensal strain of *Escherichia coli* in the guts of mice of advanced age and found that it acquired several mutations common to bacteria colonizing young mice, which were absent in old animals. This, together with the increase in *Akkermansia muciniphila* in mice of advanced age, suggest healthy aging^7,8^.

However, mutations acquired exclusively in the older were mainly pathoadaptive, tuning the metabolism to oxygen and iron availability, hypermotility, and biofilm formation.

In summary, while the evolutionary signature in the guts of very old mice shows youth-like features that may be associated with longevity, the selection of pathoadaptive traits is magnified in very old age.

While suggesting that a breach in colonization resistance is not needed to justify the abundance of age-associated pathobionts, our findings raise the question whether specialized bacteria, as opposing to generalists such as *E. coli*, will display the same ability to evolve pathoadaptive traits.

**Highlights:**

- Gut commensals face increasingly personalized selective pressures in the aging gut
- Even healthy aging selects for pathoadaptive traits in the gut microbiota
- *E. coli*‘s adaptive pattern better reflects the metabolome than microbiota composition

**In Brief:** Pathobionts are often enriched in the microbiota of the elderly. Melo-Miranda et al. showed that irrespectively of limiting the opportunity for gut invasion, the strength of selection for pathoadaptation increases with aging. Yet, the gut environment of extreme ages seems to converge, highlighting the discontinuity of the aging process.

## RESULTS AND DISCUSSION

Aging is a complex and continuous process, thus studies merely contrasting young and old individuals may overlook important age-related dynamics, resulting in an incomplete view of the process^1^. Here, we have studied the evolution of a commensal strain of *E. coli* in the guts of mice of advanced age (very old, VO, 25 months old; n=10) and compared it with old (O, 19 months old; n=6)^9^ and young (Y, 6-8 weeks old; n=15)^10,11^ animals. A measure of biological age, the frailty index^12^, showed a progressive increase with age, with significant differences between young and mice of advanced age, as expected (Fig 1A).

**Figure 1.**
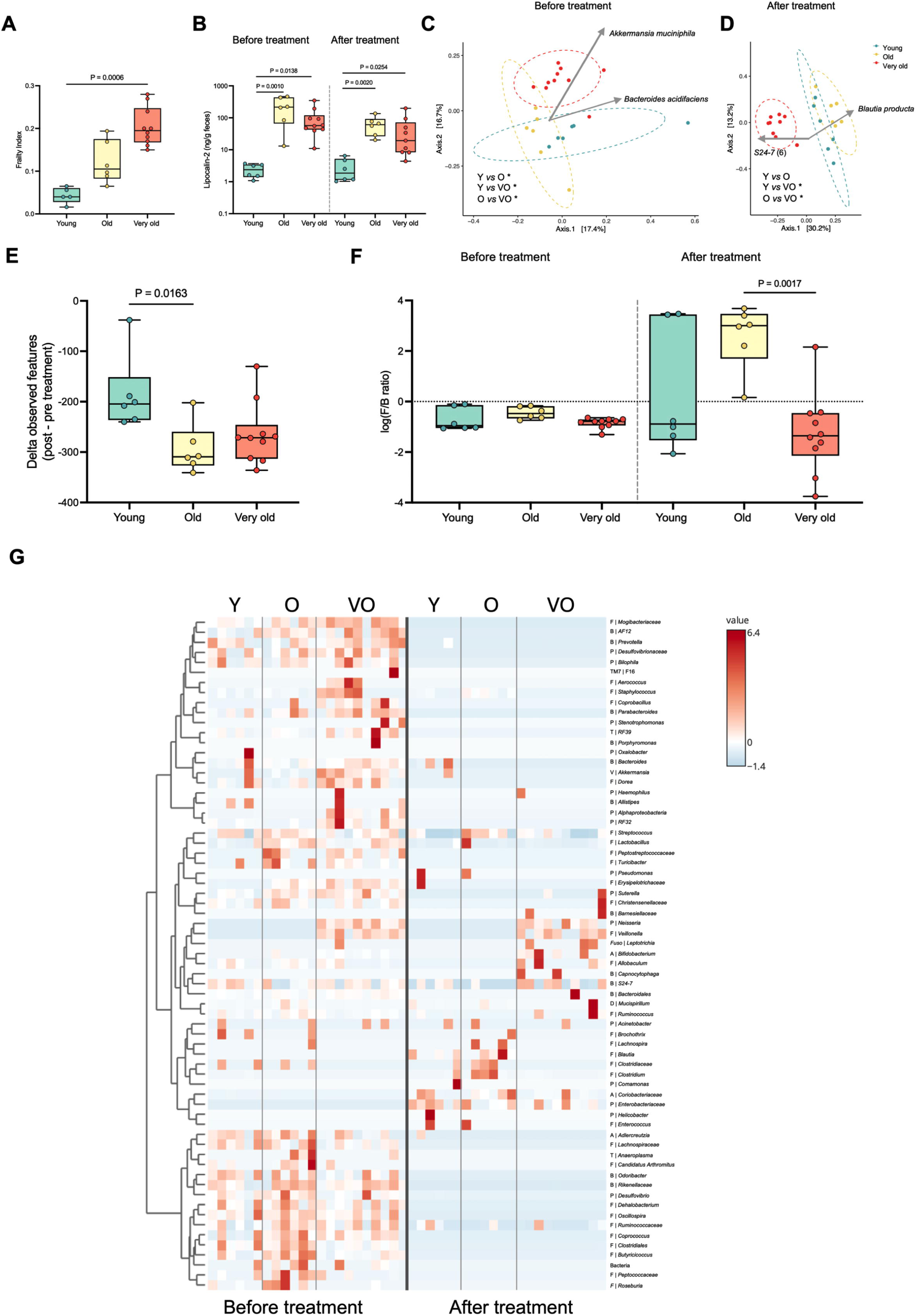
Age-Associated Changes in the Host and Gut Microbiota Composition Before and After Treatment with Streptomycin and *E. coli* Colonization. (A) Frailty index before treatment for young (n = 5)^9^, old (n = 6)^9^ and very old (n = 10) mice (Kruskal Wallis test followed by Dunn’s multiple comparison correction). (B) Intestinal inflammation, assessed via fecal lipocalin-2 levels for young (n = 5)^9^, old (n = 6)^9^, and very old (n = 9 or 10) mice, before and after treatment. Differences in the median were tested using Kruskal Wallis test followed by Dunn’s multiple comparisons correction. (C) and (D) Analysis of microbiota composition by Non-metric Multidimensional Scaling (NMDS) illustrating beta diversity of the rRNA 16S region V4 (Unweighted Unifrac) and biplots representing main contributor ASVs for young (n = 6)^9^, old (n =6)^9^, and very old (n = 10) mice, before (C) and after (D) treatment. Statistical comparisons were performed with PERMANOVA followed by pairwise post-hoc tests. * *P* < 0.05, \*\**P* < 0.01, *** *P* < 0.001, **** *P* < 0.0001. (E) Delta (after - before treatment) of the alpha diversity (observed features) of the gut microbiota of young (n = 6)^9^, old (n = 6)^9^, and very old (n = 10) mice. Differences in the median were tested using Kruskal Wallis test followed by Dunn’s multiple comparisons correction. (F) Log_10_ of Firmicutes/Bacteroidetes (F/B) ratio of the gut microbiota of young (n = 6)^9^, old (n = 6)^9^, and very old (n = 10) mice, before and after treatment. Differences in the median were tested using Kruskal Wallis test followed by Dunn’s multiple comparisons correction. (G) Heatmap illustrating the relative abundance of each OTU (genus level) of the gut microbiota of young (n = 6)^9^, old (n = 6)^9^, and very old (n = 10) mice before and after treatment. Each OTU is autoscaled (mean-centered and divided by the standard deviation of each variable). B – Bacteroidetes; D – Deferribacteres; F – Firmicutes; Fuso – Fusobacteria; P – Proteobacteria; T – Tenericutes; V – Verrucomicrobia.

We then used a previously established gut colonization system, the streptomycin-treated mouse model^10,13^, to expose age-related alterations in the intestine that could influence the evolution of their microbiota. In this colonization model^9,10^, mice were individually housed and colonized in a 1:1 proportion with two isogenic *E. coli* lineages, except for the expression of either yellow (YFP) or cyan (CFP) fluorescent proteins. These proteins serve as neutral markers, enabling the tracking of adaptive mutations in the colonizing *E. coli*. An increase in marker frequency reflects the spread of a beneficial mutation within the linked genetic background.

Successful colonization by *E. coli* requires treatment with streptomycin, which depletes multiple classes of anaerobic and facultative anaerobic bacteria^13^. Therefore, the effect of the antibiotic on age-related differences among the cohorts was assessed both before and after treatment. Inflammation, a hallmark of aging (known as inflammaging^2^), was locally evaluated by quantifying fecal levels of lipocalin-2. These were elevated in both cohorts of old mice when compared to young and remained significantly higher after treatment (Fig. 1B). The absence of further progression from old to advanced age is compatible with deceleration in inflammaging^4^.

Age-related differences in the gut microbiota composition are well-documented^14,15^ and our data confirmed that young, old and very old mice have distinct microbiota profiles (Figs. 1C, 1G, S1-S2, and Table S1). Specifically, 90% of the microbiota was composed of Bacteroidetes (∼75%), Firmicutes (10-25%) and Verrucomicrobia (0-35%) (data not shown), being significantly different between the three cohorts (Fig. 1C, PERMANOVA on Unweighted Unifrac, *P*=0.001). A particularly noteworthy distinction of mice with advanced age was the enrichment of health-associated, short-chain fatty acid (SCFA) producing-bacteria (e.g., *Akkermansia muciniphila*, an amplicon sequence variant (ASV) of *Oscillospira* sp.) (Table S1), a profile compatible with healthy aging in humans^7,8^. This cohort was further enriched in several ASVs of *S24-7* (Table S1).

As expected, streptomycin significantly reduced the microbiota diversity (Figs. 1E, 1G, S2). However, its impact was stronger in the older cohorts; a Kruskal-Wallis test on the variation (delta) in observed features (Fig. 1E) revealed a significant difference (*P*=0.012), specifically between the young and old cohorts (Dunn’s multiple comparisons test, *P*=0.0163). As a result, the microbiota composition of the advanced-age mice became even more distinct from the other two cohorts (Fig. 1D, PERMANOVA on Unweighted Unifrac, *P*=0.001). This distinction was likely driven by the differential impact of streptomycin on Bacteroidetes across the three cohorts (Fig. 1F). In the old cohort, Bacteroidetes dropped close to the detection limit in most animals (Fig. S3A-B). However, in the very old cohort, several members of the *S24-7* family (Bacteroidetes phylum) were particularly resistant to streptomycin (Figs. S3A-B). In contrast, the Firmicutes depletion by streptomycin was weaker in the old cohort than in the young and very old (Figs. S3C-D). After streptomycin, Firmicutes were mainly represented by *Lachnospiraceae*, reaching a mean frequency of ∼0.5 in the old and below 0.25 in the other two cohorts.

**Figure 2.**
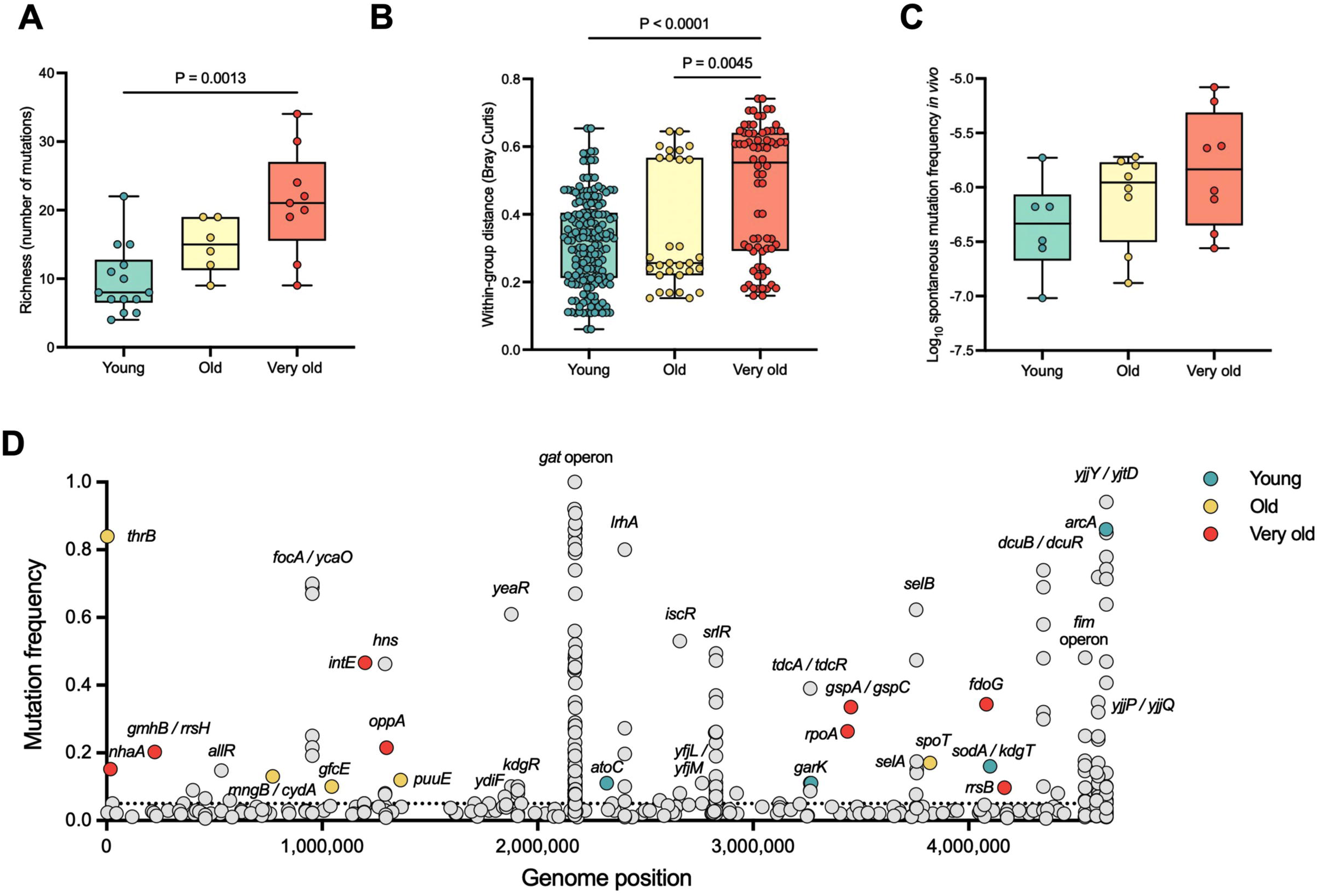
The Genomic Landscape of *E. coli* Evolution Across Ages. (A) Number of mutations acquired by *E. coli* after 24 days of evolution in the gut of young (n = 14)^9,11^, old (n = 6)^9^, and very old (n = 9) mice. Each dot represents one animal. Differences in the median were tested using Kruskal Wallis test followed by Dunn’s multiple comparisons correction. (B) Within-group distance (Bray Curtis) of the genetic diversity at day 24 of evolution among young (n = 14)^9,11^, old (n = 6)^9^, and very old (n = 9) mice. Differences in the median were tested using Kruskal Wallis test followed by Dunn’s multiple comparisons correction. (C) Spontaneous mutation frequency to furazolidone resistance in young (n = 6), old (n = 8) and very old (n = 8) mice. Differences in the median were tested using Kruskal Wallis test followed by Dunn’s multiple comparisons correction. (D) Gene targets, and its frequency and position in *E. coli*’s genome. Highlighted in color are unique gene targets mutated at a frequency > 0.1 in the young (n = 14)^9,11^, old (n = 6)^9^, and very old (n = 9) mice, at day 11 and 24 of evolution.

In summary, the very old cohort showed the biggest distinction from the three (either pre- or post-streptomycin, Fig 1C-D, respectively), but the streptomycin induced-dysbiosis was particularly visible in the old cohort, in terms of the F/B ratio (Fig. 1G). This ratio often undergoes dramatic shifts in response to antibiotic administration^16^ or metabolic disturbances (e.g., diabetes and obesity), despite contradictory results^17^

The substantial increase in *Enterobacteriaceae* after streptomycin (Fig. 1G) related to *E. coli* colonization.

### *E. coli* faces increasingly personalized selective pressures in the gut of advanced-age mice

*E. coli* colonized the gut of the two cohorts of old mice at a significantly lower density when compared to young mice (Figure S4A; group:time interaction, F(24, 322.52)=3.80, *P*<0.0001). The lower population size in the old did not result in a slower signal of adaptive evolution, as inferred from the similar dynamics of the neutral markers across age groups (Fig. S4B). However, the spread of clones with a *gat*-negative phenotype, which was previously shown to be the first adaptive phenotype in young mice^9^, was notably delayed in mice of advanced age (Fig. S4C; group:time interaction, F(10,137.08)=3.51, *P*=0.0004). This delay could be due to other mutations of greater effect driving the adaptive process of *E. coli*, and/or to the existence of a higher number of adaptive targets in the context of advanced age.

In fact, the number of segregating mutations progressively increased with age, being significantly higher in animals of advanced age compared to young animals (Fig. 2A), with a median number per animal of 21 in very old animals, but only 8 in young mice.

Additionally, the two cohorts of old animals (n=15) collectively displayed three times more unique high-frequency targets (frequency >0.1) than young animals, with 12 targets in old versus only 4 in young animals (Fig. 2D, Table S3, and ^9,11^), explaining the heterogeneity and diversity increase in segregating mutations (Figs. 2B, S5).

Since the number of segregating mutations is influenced by both their fitness effect and rate of production, the frequency of furazolidone resistant clones in feces was measured as an estimate of the spontaneous mutation rate in the gut. A progressive increase in the *E. coli*’s spontaneous mutation rate with age was observed (Fig. 2C), albeit not reaching statistical significance.

### Pathoadaptive traits spread in the gut microbiota with advanced age, even under healthy conditions

Changes in selective pressures faced by *E. coli* colonizing the aged gut were identified by focusing on parallel targets of mutation (i.e., mutations occurring in two or more mice, Tables S2-S4), as parallelism is a signature of adaptation^18^. We further queried whether those could be related to the common observation of pathobiont enrichment in old age^4^.

Comparing parallel with cohort-specific mutations can help disentangle age from environmental-related adaptations.

Mutations occurring in the *gat* operon, in the *srlR* and *hns* genes, and in the *tdcA*/*tdcR* intergenic region, were consistently observed in *E. coli* evolving in the three groups (Figs. 2D, 3A, S6, Tables S3-S4), and were deemed related to the gut environment. Most of these targets are metabolism-associated (Table S4), reflecting optimizations for efficient nutrient consumption^19,20^, irrespective of age.

**Figure 3.**
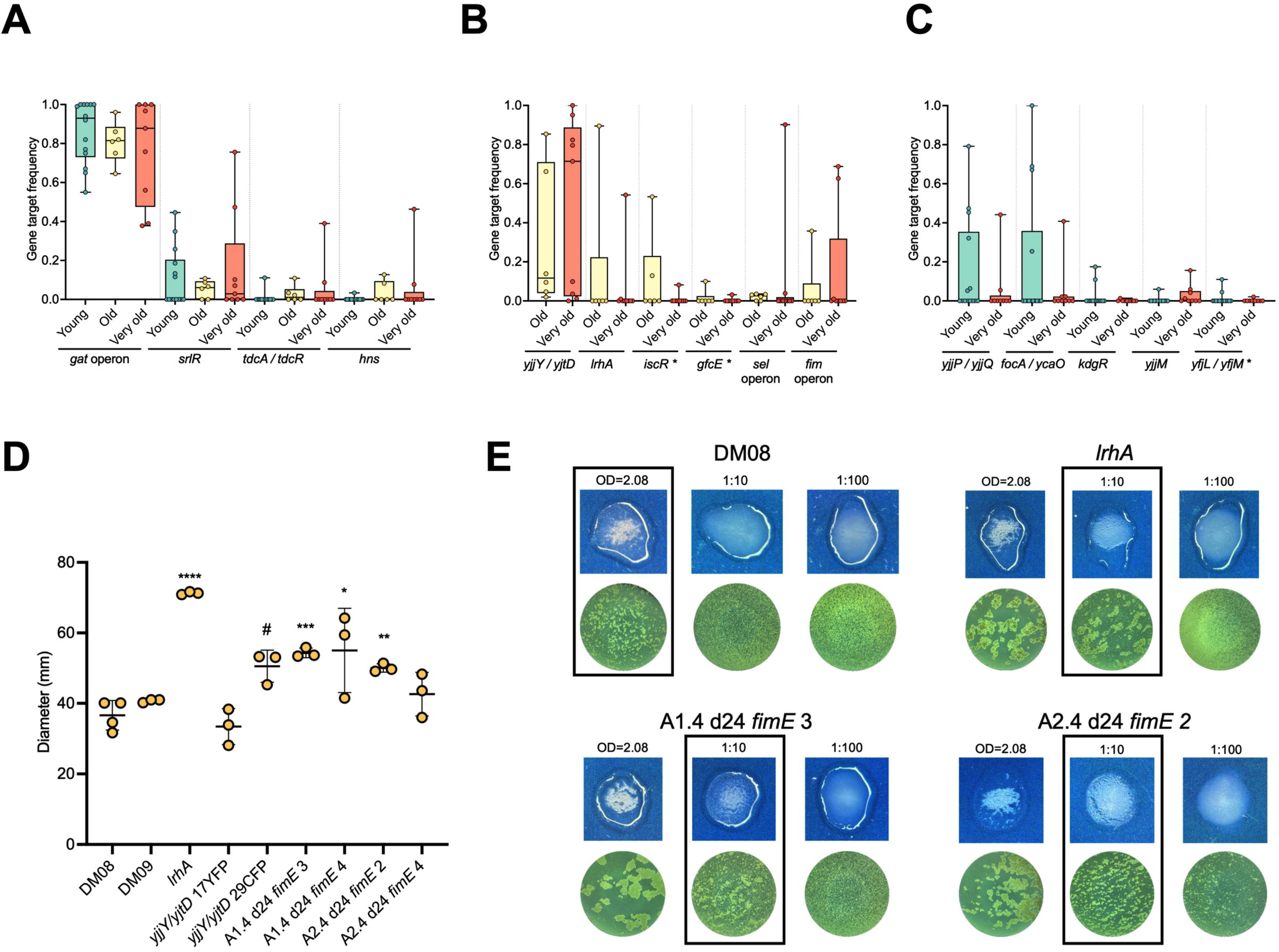
The Aged Gut Selects for Pathoadaptations, Including Hypermotility and Increased Adhesion. (A) Frequency of the mutational targets per mouse shared between young (n = 14)^9,11^, old (n = 6)^9^, and very old (n = 9) mice at day 24 of evolution. * – mutations found at day 11 of evolution in very old mice. (B) Frequency of the mutational targets per mouse shared between old (n = 6)^9^ and very old (n = 9) mice at day 24 of evolution. (C) Frequency of the mutational targets per mouse exclusively shared between young (n = 14)^9,11^ and very old (n = 9) mice at day 24 of evolution. * – mutations found at day 11 of evolution in very old mice. (D) Swarming motility of the ancestral colonizing strains (DM08 and DM09) and of the evolved clones carrying mutations in *lrhA*, *yjjY* / *yjtD* or *fimE* clones. The quantification reflects the measurement of the diameter of growth (n= 6 technical replicates). Unpaired t-test was used to compare each mutant with the ancestral carrying the same neutral marker (YFP or CFP). * – comparison with DM08; # – comparison with DM09. (E) Representative images (top, in blue: macroscopic image; bottom, in green: microscopic image) from the aggregates formed by interaction of *E. coli* with yeast. Top left panel: ancestral strain; top right panel: *lrhA* evolved mutant; bottom panels: 2 of the *fimE* clones evolved in different mice. This assay was performed in 2 independent replicates.

We then focused on targets exclusively shared among old animals (Figs. 3B, S6). Six targets were identified, from which *iscR, lrhA, fim* operon and *yjjY*/ *yjtD* were related to stress responses (Table S4) and commonly associated with pathoadaptation^21–23^. *iscR* encodes an iron-sulfur cluster regulator, one of the major regulators of iron homeostasis^24^. Similarly to ^9^, the observed mutation in this gene (Table S2) is predicted to disrupt the coordination of the iron-sulfur cluster^25^, slightly increasing *sufA* expression; in turn, the *sufABCDSE* operon encodes the machinery to build essential Fe-S clusters under oxidative stress and iron starvation^24^.

*lrhA* is a global regulator reported to repress RpoS, involved in stress responses^26^, but also in motility, chemotaxis, flagellum synthesis^27^, and type 1 fimbriae expression^28^. Observed mutations in this work and before^9^ lead to increased swarming motility (Figs. 3D and S7A), likely due to gene inactivation.

The selection of hypermotility could enhance stable colonization of the gut by *E. coli*, similar to what has been shown in *Lactobacillus agili*s, where the motile strain was able to better penetrate the mucus layer when compared to a non-motile strain^29^. In addition, hypermotility has also been associated with inflammation^30^.

The *fim* operon is involved in the production of type 1 fimbriae, essential for adhesion and involved in biofilm formation^31^. In particular, FimE is a recombinase that catalyzes the inversion of a DNA element, the *fim* switch (*fimS*), which contains the promoter for the expression of fimbrial genes. FimE is responsible mainly for ON (fimbrial gene expression) to OFF (no expression) switching^32^.

The mutations observed during evolution in the murine gut occurred in the intergenic region between *fimE* / *fimA* or in the coding region of *fimE* gene (e.g. insertion of insertion sequences (IS); Table S3). These mutations, especially IS insertions within *fimE*, suggest gene inactivation, leading to a predominantly ON phenotype, which could enhance the expression of type 1 fimbriae and biofilm formation To test this, *fimE* mutants isolated from the very old cohort, as well as a *lrhA* mutant, were tested for increased aggregation using a yeast aggregation assay. Aggregation serves as evidence of type 1 fimbriae expression^33^. As predicted, *lrhA* and *fimE* mutants demonstrated at least 10-fold increase in the ability to aggregate yeast compared to the ancestor (Figs. 3E and S7B), strongly indicating a higher ability to form biofilms. An enhanced biofilm capacity has been associated with immune system evasion, antibiotic resistance, and pathogenicity in *E. coli*^34,35^.

The *fim* operon has also been implicated in motility^36^. In agreement, three out of four *fimE* mutants (isolated from two animals) exhibited significantly higher swarming motility when compared to the ancestor (Figs. 3D and S7A).

The fourth target specific to the aged intestine was the intergenic region between *yjjY*/*yjtD.* These were previously shown to upregulate *arcA*^37^, a global regulator predicted to control the expression of genes involved in transitioning from aerobic to anaerobic respiration^38^ and *fliA*, a small sigma factor necessary for flagellum synthesis and swarming motility^39^. One of two *yjjY/yjtD* mutants tested for swarming motility showed increased motility whereas the other remained undistinguishable from the ancestor (Figs. 3D and S7A). This is consistent with the hypothesis that some of these highly prevalent mutants (present in all but one animal in the aged cohorts) are also selected for increased motility, a trait that appears pivotal in the aged gut.

Together, these targets show a signature of pathoadaptation in aged animals, namely adaptation to iron scarcity and oxygen availability, selection for hypermotility, and biofilm formation.

We then examined common targets between young and very old mice. Strikingly, five mutational targets were unique to young and advanced-age animals, contrasting with only one target exclusive to both young and old mice (Figs. 3C and S6). Interestingly, this discontinuity during aging has been observed in other contexts, such as the association between DNA methylation and aging^40^. Another feature where similarities between centenarians and young individuals were observed and linked to longevity is the microbiota composition^15^.

In our study, most targets mutated exclusively in young and advanced-age animals were associated with metabolic adaptations (Table S4), as is the case for *yjjP*/*yjjQ* (succinate transport^41^), *focA*/*ycaO* (formate metabolism^42^), *kdgR* (D-glucuronate metabolism^43^), and *yjjM* (L-galactonate consumption^44^). These suggested that, for a gut commensal such as *E. coli*, the gut metabolic environment of the advanced age is closer to the young than to that of old animals. In agreement, the ratio between metabolism over stress-related mutations was 2.5 and 1.2 in young and advanced age, respectively, versus 0.8 in the old cohort.

### The metabolic environment better relates with *E. coli* adaptive pattern than with the microbiota composition

To further characterize the metabolic environment of the intestinal tract, an untargeted metabolomics analysis using Proton nuclear magnetic resonance (^1^H-NMR) on fecal samples from all age cohorts was performed.

The metabolomes were clearly distinct between age groups, both before and after streptomycin treatment and colonization with *E. coli* (Figs. 4A, PERMANOVA *P*=0.001, and 4B, PERMANOVA *P*=0.001). This separation was further corroborated by an unsupervised clustering analysis, which correctly placed 83% of the individuals in three distinct clusters (Figs. 4C and 4D).

**Figure 4.**
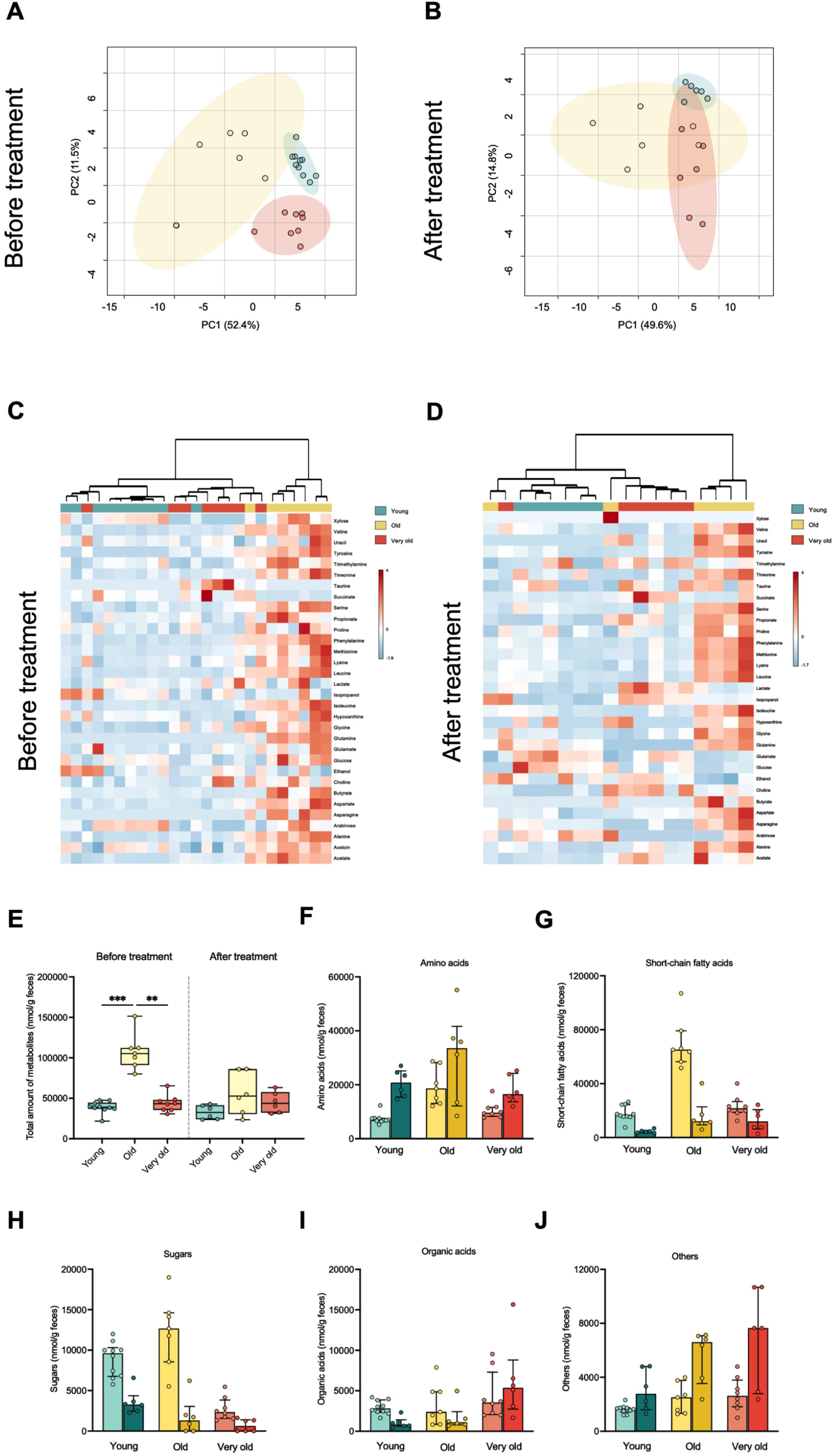
The Gut Metabolic Profile of Young, Old and Very Old Mice Before and After Streptomycin Treatment and Colonization. (A) Principal component analysis (PCA) and (C) heatmap showing distinct metabolic profiles in fecal samples of young (Y, n = 10), old (O, n = 6), and very old (VO, n = 8) mice before treatment. Statistical comparisons in the PCA plots were performed with PERMANOVA followed by pairwise post-hoc tests. In the heatmaps, each metabolite is autoscaled (mean-centered and divided by the standard deviation of each variable). The clustering analysis is unsupervised and based on Euclidean distances. (B) PCA and (D) heatmap showing distinct metabolic profiles in fecal samples of young (Y, n = 6), old (O, n = 6), and very old (VO, n = 6) mice before (A and B) and after (C and D) treatment. Statistical comparisons in the PCA plots were performed with PERMANOVA followed by pairwise post-hoc tests. In the heatmaps, each metabolite is autoscaled (mean-centered and divided by the standard deviation of each variable). The clustering analysis is unsupervised and based on Euclidean distances. (E) Total amount of metabolites (nmol/g feces) measured in the fecal samples of young, old, and very old mice, both before and after treatment. Boxplots show the median, interquartile range, and individual data points. Statistical comparisons were made with Kruskal-Wallis test followed by Dunn’s multiple comparisons test. (F-J) Bar plots showing the median concentrations of various metabolite categories before (lighter colors) and after treatment (darker colors) across the different age groups. Barplots show the median, interquartile range, and individual data points. (F) Amino acids; (G) Short-chain fatty acids; (H) Sugars; (I) Organic acids; (J) Other metabolites.

The diversity and abundance of metabolites in feces reflects the host’s absorption efficiency and the microbiota’s capacity to produce and break down these molecules. Here we observed that the total amount of the identified metabolites increases with aging, though the trend is reversed for amino acids, short chain fatty acids (SCFA) and sugars in the very old cohort (Figs. 4E-J, S8). Nutrient malabsorption is commonly reported in the elderly^45^ and could explain the observed increase in the older cohort. However, the reversal of this effect in the very old is more challenging to explain and warrants further investigation.

Considering global metabolic categories, streptomycin treatment and colonization led to an increase in amino acid concentrations and a reduction in SCFAs and sugars in an age-independent manner (Figs. 4F–H and S8A-F), indicating that colonization-induced changes did not override age-related changes in the fecal metabolome.

Importantly, heatmaps of the identified metabolites highlighted that both before and after colonization, old mice clustered apart from young and very old mice (Figs. 4C-D), driven by higher levels of total metabolites, specifically the majority of the amino acids, acetate, butyrate and trimethylamine (Figs. 4E, S8A-D, I-J). This contrasts with a study on young and old male BALB/c mice, where a decline in amino acids like isoleucine and methionine was reported in old mice^46^. Interestingly, our results are more aligned with observations of elevated levels of amino acids in a mouse model of inflammatory bowel disease (IBD) colonized with *E. coli*^47^ and branched-chain amino acids (leucine, isoleucine, and valine)^48^ in individuals with IBD, although the levels of intestinal inflammation we observed in the old cohort are likely lower than those observed in pathological conditions. In the context of IBD, this may occur due to malabsorption^49^.

SCFAs are largely produced by bacteria and are generally associated with positive health outcomes^50^. In humans, previous studies have shown that fecal SCFA levels decrease with age^51^, while mice studies did not find significant differences between young and old^46^. Intriguingly, old mice showed the highest levels of most SCFAs, both before and after colonization (Fig. 4G, S8C-D). This could be related with the butyrate producer *Blautia producta* (family *Lachnospiraceae*)^52^, which was boosted in the old cohort after antibiotic (Fig 1D and Fig. S3D).

The advanced-aged cohort displayed unique features. Specifically, the concentrations of arabinose, glucose and xylose were consistently lower in very old mice than in young and old animals, with xylose becoming undetectable in any group post-colonization (Fig. S8E-F). Additionally, prior to antibiotic treatment and colonization, taurine was only found in feces of very old mice. Taurine in circulation typically decreases with age, and taurine supplementation has been shown to increase healthspan and lifespan in various model organisms^53^. Our observation raises two possibilities: either this is an indication of a physiological trait linked to healthy aging, or it reflects impaired taurine absorption, accumulating in feces. While nutrient absorption generally declines with age, we did not observe such a trend for any of the other metabolites (Figs. S8).

Overall, the metabolic environment encountered by *E. coli* during colonization was more similar in animals at the extremes of the age range — the youngest and the oldest, aligning with our findings on the adaptive patterns of *E. coli*.

## Conclusion

Aging is not linear and, therefore, studies encompassing multiple timepoints are essential to fully describe its dynamics. Here, we queried changes in the microbiome environment across aging that can contribute to pathobiont enrichment in the elderly, including cases of healthy aging.

Even under the controlled conditions of our study – standardized diet, environmental stability, and stringent measures to prevent colonization by pathogens – we observed the emergence of pathoadaptations in old age. This strongly suggests that the aging process provides an internal environment prone for the selection of potentially pathogenic traits within the commensal gut microbiota.

While the composition of the microbiota certainly shapes the metabolomic landscape^54^, it appears that *E. coli* evolution more effectively captures differences between age-related metabolic environments. This finding is unexpected, given that although *E. coli* constitutes a significant portion of the microbiota across all age groups after colonization, it represents a single strain of a single species. One would expect that differences in its adaptive response would reflect its own specific needs, while global changes in microbiota composition would reveal more general properties of the metabolome. However, us and others^55^ have shown that experimental evolution using *E. coli,* a genetically versatile and metabolically flexible gut commensal, is a powerful strategy to describe changes in the gut environment.

Our work highlights the impact that aging can have on the gut microbiota, selecting for pathoadaptive traits. Future work should address if more specialized bacteria will be able to respond so efficiently as *E. coli* to age-associated gut changes.

Future studies could focus at exploring whether interventions aiming at delaying or attenuating aging (e.g. caloric restriction or exercise) could also act on intestinal aging and subsequently mitigate the selection of pathoadaptive traits.

## ACKNOWLEDGMENTS

We thank Marta Teixeira Pinto from Instituto de Investigação e Inovação em Saúde (i3S) for allowing the use of the fluorescent stereoscope which enabled the quantification of DM08-YFP and and DM09-CFP *E. coli*.

This work was funded by Fundação para a Ciência e Tecnologia PTDC/BIAEVL/30212/2017 (“MicroAgeing – the role of the microbiota in ageing”), and Programa Operacional Regional do Centro, through Fundo Europeu de Desenvolvimento Regional—FEDER (CENTRO-01-0145-FEDER-030212), and by FCT - Fundação para a Ciência e Tecnologia, I.P. by project reference UIDB/04501/2020 and DOI identifier https://doi.org/10.54499/UIDB/04501/2020 and project reference UIDP/04501/2020 and DOI identifier https://doi.org/10.54499/UIDP/04501/2020.

R. M-M. was supported by the individual Grant 2020.05130.BD (DOI identifier: https://doi.org/10.54499/2020.05130.BD) from FCT.

H.C.B. was supported by the DREAM ANR-20-AMR-0002 grant, and by the HORIZON-MSCA-2023-PF-01 project number 101148351 - MICROINVADER, funded by the European Union. Views and opinions expressed are however those of the author only and do not necessarily reflect those of the European Union or the European Research Executive Agency.

This work was also supported by CICECO – Aveiro Institute of Materials (UIDB/50011/2020, DOI 10.54499/UIDB/50011/2020; UIDP/50011/2020, DOI 10.54499/UIDP/50011/2020 & LA/P/0006/2020, DOI 10.54499/LA/P/0006/2020), and LAQV-REQUIMTE (LA/P/0008/202, DOI 10.54499/LA/P/0008/2020; UIDP/50006/2020, DOI 10.54499/UIDP/50006/2020; UIDB/50006/2020, DOI10.54499/UIDB/50006/2020), financed through national funds through FCT/MCTES. FCT is also acknowledged for the research contract under the Scientific Employment Stimulus to I.F.D. (CEECIND/02387/2018). The NMR spectrometer is part of the National NMR Network (PTNMR) and is partially supported by Infrastructure Project N° 022161.

A. S. was funded by FCT through Estímulo Ao Emprego Científico (reference CEECINST/00026/2018/CP1521/CT0001 and DOI identifier https://doi.org/10.54499/CEECINST/00026/2018/CP1521/CT0001).

## AUTHOR CONTRIBUTIONS

Conceptualization, R.M-M. and A.S.; methodology, R.M-M. and A.S.; validation, R.M-M. and A.S.; formal analysis, R.M-M., C.J. and I.D.; investigation, R.M-M. and A.P.; resources, A.S.; writing – original draft, R.M-M. and A.S..; writing – review & editing, R.M-M., A.S., H.C.B., I.G., I.D. and A.P.; visualization, R.M-M. and A.S.; funding acquisition and supervision, A.S.

## STAR METHODS

### RESOURCE AVAILABILITY

#### Lead contact

Further information and requests for resources and reagents should be directed to and will be fulfilled by the lead contact, Ana Sousa.

#### Materials availability

This study did not generate new unique reagents.

#### Data and code availability

- This paper analyzes existing, publicly available data (genome sequencing data of old and very old mice), under the accession number PRJNA588787 in the NCBI BioProject database (https://www.ncbi.nlm.nih.gov/bioproject/).
- 16S rRNA gene sequencing data (for all experimental groups) and genome sequencing data (for the very old group) have been deposited under the BioProject accession number XXXXX in the NCBI BioProject database (https://www.ncbi.nlm.nih.gov/bioproject/).
- This paper does not report original code.
- Any additional information required to reanalyze the data reported in this paper is available from the lead contact upon request.

### EXPERIMENTAL MODEL AND STUDY PARTICIPANT DETAILS

#### Mice

Female C57BL6/6J very old mice (25 months old) were reared under specific pathogen-free barrier conditions at the Instituto Gulbenkian Institute for Molecular Medicine (GIMM) animal facility at Oeiras Campus. The animals were then transferred and single housed in individually ventilated cages with access to food and water *ad libitum* at Instituto de Biomedicina (iBiMED) animal facility. Animals were allowed to acclimate for at least 1 week before any experimental procedure.

This research project was ethically reviewed and approved by the Animal Welfare Body of the iBiMED (license reference: B003/2021), and by the Portuguese National Entity that regulates the use of laboratory animals (DGAV – Direção Geral de Alimentação e Veterinária (license reference: 0421/000/000/2021).

Young (6-8 weeks) and old (19 months) mice data originated from Barroso-Batista et al. and Barreto et al^9–11^. Exceptions are the animals used to perform the spontaneous mutation rate and metabolomics experiments, where another set of young (6-8 weeks) female mice from the animal house of GIMM were used (license reference: A009.2018; DGAV license reference: 009676).

All experiments conducted on animals followed the Portuguese (Decreto-Lei no 113/2013) and European (Directive 2010/63/EU) legislations, concerning housing, husbandry and animal welfare.

#### Bacteria

All bacterial strains utilized in this study were derived from *E. coli* K-12 MG1655. The strains DM08-YFP and DM09-CFP carry either the yellow (YFP) or cyan (CFP) fluorescent genes, respectively, which are integrated into the *ampR* locus at the *galK* locus and are constitutively expressed. These strains also exhibit streptomycin resistance due to a K43R amino acid substitution in the *rpsL* gene. Both strains were previously engineered and documented in a previous evolution study^10^. Refer to the Key Resources Table for a comprehensive list of all strains and primers employed in this work. *E. coli* cultures were routinely propagated in Lysogeny Broth (LB, Lennox formulation) at 37 °C with aeration, unless otherwise specified.

### METHOD DETAILS

#### Measurement of intestinal inflammation

To assess intestinal inflammation through fecal lipocalin-2 concentration, fresh fecal pellets were collected and immediately frozen at -80 °C without any buffer. Subsequently, the samples were homogenized in PBS 1X to achieve a final concentration of 100 mg/mL. After vortexing for 5 minutes, the homogenates were centrifuged at 18000 g for 15 minutes at 4 °C. The resulting supernatant was collected and stored at -20 °C until further analysis. Lipocalin-2 concentration was determined using ELISA following the manufacturer’s instructions (Mouse lipocalin-2/NGAL DuoSet ELISA, R&D Systems), with a dilution of 1:10 for most samples.

#### Microbiota analysis

To describe the bacterial context in which *E. coli* evolved, we re-sequenced the frozen fecal samples collected from young and old animals^9^ and sequenced samples from very old mice before and during the evolution experiment. The DNA extraction was performed using QIAamp DNA Stool Mini Kit (QIAGEN), according to the manufacturer’s instructions plus a step of mechanical disruption with glass beads (0.1 mm) and shaking at room temperature, 1500 rpm, for 30 minutes. Sequencing of the V4 region of 16S rRNA gene (primers 515F-806R) was performed at the Genomics Unit of GIMM, employing a 280-multiplex approach on a 2x250bp PE MiSeq run. Raw reads were processed using QIIME2 2020.8 with default parameters^56^. Deblur was used for quality filtering, denoising, and ASV calling^57^.

The R package *decontam* with the prevalence method of contaminant identification^58^ was used to exclude ASVs determined as contaminants from the samples, based on the microbiota composition of 14 negative controls.

The resulting filtered ASV table was then used in the R package *phyloseq*^59^ to determine beta-diversity metrics, using rarefaction based on the size of the sample with least sequencing depth. Alpha diversity was calculated on QIIME2 with *diversity core-metrics-phylogenetic* and *diversity alpha-group significance*, again with rarefaction value based on the sample with the lowest coverage.

Differentially abundant ASVs between young, old, and very old animals were identified using ANCOM 2.0^60^ with a W cut-off significance threshold of 0.8. For ANCOM, *exactRankTests*, *nlme*^61^ and *gglplot2*^62^ packages were used.

Taxonomical classification was determined by matching ASVs against the Greengenes 13.8 database with *feature-classifier classify-sklearn*, with further validation conducted via NCBI.

#### *E. coli* colonization

To track the evolutionary trajectory of *E. coli* in the gut of very old mice, we used the classical streptomycin-treated mouse colonization model as described before ^10,13^. In brief, mice were given autoclaved water containing streptomycin (5 g/L) for 24 hours. Following a period of 4 h of water and food deprivation, mice were orally gavaged with 100 µL of a 10^8^ suspension of a 1:1 mixture of the *E. coli* strains DM08-YFP/DM09-CFP in PBS 1X. After gavage, animals had the streptomycin (5 g/L) supplemented water and food returned. Throughout the experiment, the streptomycin supplemented water was replenished every 3 days. Fecal samples were collected over a 24-day period, homogenized in PBS 1X and plated in LB agar supplemented with streptomycin (100 µg/mL) to selectively culture *E. coli* strains, thus following their evolution along time, or stored with 15% glycerol at -80 °C for future analysis. Appropriate dilutions were then plated and incubated overnight at 37°C in aerobiosis. Total bacterial loads and the frequency of YFP and CFP bacteria was assessed using a fluorescent stereomicroscope (Olympus Stereomicroscope SZX16). The evolutionary trajectory of *E. coli* in the gut of young and old mice were explored in previous studies following the same experimental design ^9–11^.

#### Dynamics of the *gat*-negative phenotype

The frequency of the *gat*-negative phenotype throughout the experimental evolution in young and old mice was determined in ^9–11^.

To monitor the same phenotype in very old mice, we employed the same previously described method^10^. Briefly, frozen fecal pellets were serially diluted to the appropriate concentrations and plated onto MacConkey agar supplemented with 0.4% galactitol (Dulcitol). Following overnight incubation at 37°C, we were able to differentiate between bacteria capable of metabolizing galactitol (yielding red colonies) and those unable to do so (resulting in white colonies). The frequency of mutants was determined by calculating the ratio of white colonies (indicative of gat-negative bacteria) to the total number of colonies.

#### Whole genome re-sequencing

Data for both young and old mice were obtained from ^9,11^.

For very old mice, frozen fecal pellets underwent serial dilution and culture on LB agar plates supplemented with streptomycin (100 μg/mL). After overnight incubation at 37 °C, a mixture of >1000 clones was collected from the plates and resuspended in 10 mL PBS 1X. DNA extraction was performed using with phenol-chloroform^63^. Library construction (Pico Nextera) and sequencing were conducted at the GIMM Genomics Unit using an Illumina NextSeq 500 Sequencer. Each sample was pair-end sequenced, generating datasets of paired-end 250 bp read pairs.

Sequencing adapters were automatically detected and removed using fastp v0.23.2^64^. Raw reads were trimmed from both ends, using window sizes of 4 base pairs ensuring an average base quality of a minimum value of 30 to be retained. Trimmed reads were kept if they met a minimum length of 100 bps and consisted of at least 50% base pairs with phred scores of at least 20.

Then, optical duplicates known to occur in NextSeq sequencing were removed using clumpify from the BBTools package^65^.

Mutations were called using the 0.23 version of the BRESEQ pipeline^66^ with polymorphism option on and default settings except for i) requirement of a minimum coverage of 3 reads on each strand per polymorphism; ii) eliminating polymorphism predictions occurring in homopolymers of length greater than 3; iii) discarding polymorphism predictions with significant (p < 0.05) strand or base quality score bias; iv) polymorphism frequency cutoff of 0.

All mutations were manually curated using Integrative Genome Viewer (IGV)^67^.

The Venn Diagram of the mutational targets was done using *VennDiagram*^68^ R package.

#### Mutational diversity analysis

We performed an analysis like the one described by Turner et al.^69^, as previously employed in^9^. This approach uses the Bray-Curtis dissimilarity index to gauge genetic similarities between populations. Specifically, the R package *vegan*^70^ was used to perform a non-metric multi-dimensional scaling (NMDS) using the Bray-Curtis dissimilarity index, considering the frequency of the mutations in each population. In instances where certain genes exhibited multiple mutations within the same population, the sum of those different mutations was used. Mutations in the *gat* operon were treated as mutations in a single gene, given their shared phenotype (the loss of ability to consume galactitol). Permutational analysis of variance (PERMANOVA) was then conducted to compare the mutational diversity of the three age groups, with significance determined at a p-value threshold below 0.05. The within-group distance as a measure of group heterogeneity was calculated using the Bray-Curtis dissimilarity matrix.

#### Mutation rate *in vivo*

Mutations that confer furazolidone resistance are expected to be found in *E. coli* populations at a mutation-selection balance and their frequency to reach a stable equilibrium, which is proportional to the mutation rate of the genome^55^.

To estimate frequency of furazolidone resistance clones, we determined the fraction of E. coli clones carrying spontaneous resistance to furazolidone, as previously described^11,55^. In summary, fecal samples from day 24 of experimental evolution preserved in 15% glycerol were plated on LB agar supplemented with streptomycin (100 µg/mL) in the appropriate dilutions to count the total number of *E. coli* CFUs, or LB agar supplemented with streptomycin (100 µg/mL) and furazolidone (1.25 µg/mL) to count the number of *E. coli* CFUs that were resistant to furazolidone. After incubation overnight at 37 °C, plates were counted and the frequency of mutants resistant to furazolidone was calculated as the ratio between the number of furazolidone-resistant clones and the total loads in each *E. coli* population.

#### Isolation of *fimE* mutants

Two *fimE* mutants per mouse, carrying either IS2 (1331 bp) and IS5 (1195 bp) insertions, were isolated from the *E. coli* population of mice A1.4 and A2.4 at day 24 of evolution. To do so, fecal samples were inoculated and grown overnight at 37 °C in LB supplemented with streptomycin (100 µg/mL). Random colonies (14 per mouse) were screened. Since the *fimE* mutation was due to an IS insertion in the coding region, the *fimE* gene size of regular clones is 668 bp and the size in mutants is of 2009 bp or 1871 bp (depending on whether it is the IS2 or IS5 insertion, respectively). So, we performed a PCR to amplify *fimE*, using fimE_Fw and fimE_R primers (see Key Resources Table).

The PCR reaction was performed in a total volume of 50 µL, containing directly picked bacterial culture, 1X Taq polymerase buffer, 0.25 mM dNTPs, 2.5 mM MgCl_2_, 0.25 µM of each primer, and 1.25 U of Taq polymerase. PCR conditions were as follows: 95°C for 5 min, followed by 30 cycles of 95°C for 30 s, 60°C for 30 s, and 72°C for 31 min, finalizing with 5 min at 72 °C.

#### Swarming motility

For the swarming motility assay, three independent colonies of the ancestral strains DM08 and DM09, as well as the *yjjY* / *yjtD* mutants (isolated in ^37^), the *lrhA* mutant (isolated in ^9^) and the four isolated *fimE* mutants were grown overnight in LB at 37 °C with agitation and aeration. 3 µL of the overnight culture were then used to inoculate the center of a 25 mL-thick Tryptone Swarming Agar plate (5 g/L Tryptone, 2.5 g/L NaCl, 1.25 g/L Agar) ^71^. Bacteria were incubated for 18 h at 37 °C. A minimum of 6 plates were inoculated for each clone, and the experiment was independently performed at least three times. Photographs of the plates were captured using GelDoc XR+ (Bio-Rad) under white light, and diameter measurements were conducted using ImageJ/Fiji^72^.

#### Yeast agglutination assay

For the yeast agglutination assay, the ancestral DM08, the *lrhA* mutant ^9^ and the four isolated *fimE* mutants were grown overnight in LB at 37 °C with agitation and aeration. Then, optical density at 600 nm (OD600) was measured and normalized to an OD of 2, from which 10^-1^ and 10^-2^ dilutions were performed. 10 µL of a fresh solution of yeast powder (10 mg/mL of lyophilized yeast powder in 1x PBS) was mixed with 10 µL of each bacterial dilution and dropped onto a microscope glass slide (adapted from^28^). The drops were allowed to settle undisturbed for 10 minutes. The entire drop was observed and photographed using a stereo microscope and an optical microscope.

#### Metabolomics of fecal samples

We performed ^1^H-NMR analysis of mouse fecal samples to determine the composition of the metabolic environment in the mouse gut of the three age cohorts, using a protocol adapted from ^55^. In brief, fecal samples (70 mg) were diluted in 700 µL of phosphate buffer (PBS), transferred into microtubes containing 0.3 g of 0.1 mm glass beads (Scientific Industries SI-BG01) and bead-beaten using Mixer Mill MM 400 (Retsch), 2X1 min with 30 rev/s pulses. The samples were then centrifuged at 20817 g for 20 min at 4 °C to remove debris and the glass beads. Supernatants were collected and filtered through 0.22 µm filters (Milipore). Next, samples were ultra-filtered through pre-rinsed 3 KDa spin columns (Vivaspin 500) by centrifugation at 15000 g and 4°C, for 3 h or until 150 µL of filtrate was obtained. Filtered samples were then dried under vacuum (Centrivap, model 73100, Labconco, Kansas City, MO, USA), and the extracts stored at -80 °C until spectral acquisition.

For spectral acquisition, extracts were reconstituted in 550 µL of deuterated phosphate buffer (PBS 100 mM, pH 7), containing 0.1 mM of 3-(trimethylsilyl)propionic acid (TSP-d_4_). Each sample was transferred to a 5 mm NMR glass tube.

Spectra were acquired on a Bruker Avance III HD 500 NMR spectrometer (University of Aveiro, Portuguese NMR Network) operating at 500.13 MHz for ^1^H observation, at 298K. One-dimensional (1D) ^1^H spectra were recorded with 32 k points, 7002.80 Hz spectral width, a 2 s relaxation delay, and 512 scans, using the pulse programs “noesypr1d”. These spectra were processed in TopSpin 4.0.3 (Bruker BioSpin, Rheinstetten, Germany), using exponential multiplication (LB 0.3 Hz), zero-filling to 64 k data points, manual phasing, baseline correction, and calibration to the TSP-d_4_ (0 ppm). Two-dimensional (2D) NMR spectra, namely ^1^H-^1^H TOCSY, *J*-resolved and ^1^H-^13^C HSQC spectra, were also recorded for selected samples to aid metabolite identification. Metabolite assignment was based on matching 1D and 2D spectral information to reference spectra available in Chenomx (Edmonton, Canada) and BBIOREFCODE-2–0–0 (Bruker Biospin, Rheinstetten, Germany). Metabolite quantification was performed using signal deconvolution and fitting to reference spectra available in the Chenomx library through the Profiler module of the Chenomx NMR Suite V10.1 Professional.

All the remaining analyses were performed with Metaboanalyst 6.0^73^.

### QUANTIFICATION AND STATISTICAL ANALYSIS

All statistical analyses were performed using GraphPad Prism 9 for Mac (GraphPad Software, San Diego, California, USA, www.graphpad.com) or the statistical software R version 4.2.1^74^. Detailed statistical information for each experiment is provided in the figure legends and/or within the manuscript, including sample sizes. In figures, *n* corresponds to the number of animals, except for Fig. 3B (swarming motility), where *n* represents the number of biological independent replicates performed.

Before applying the Unpaired t-test or the linear mixed models, the normality of data distributions was assessed using the Shapiro-Wilk normality test and QQ plot visualization.

Statistical significance was defined as *p* < 0.05 for all comparisons and was calculated according to the methods outlined in the manuscript and/or figure legends. * *P* < 0.05; ** *P* < 0.01; *** *P* < 0.001; **** *P* < 0.0001.

