## Supplemental Figures and Tables for "Aging: an inevitable road toward gut microbiota pathoadaptation"

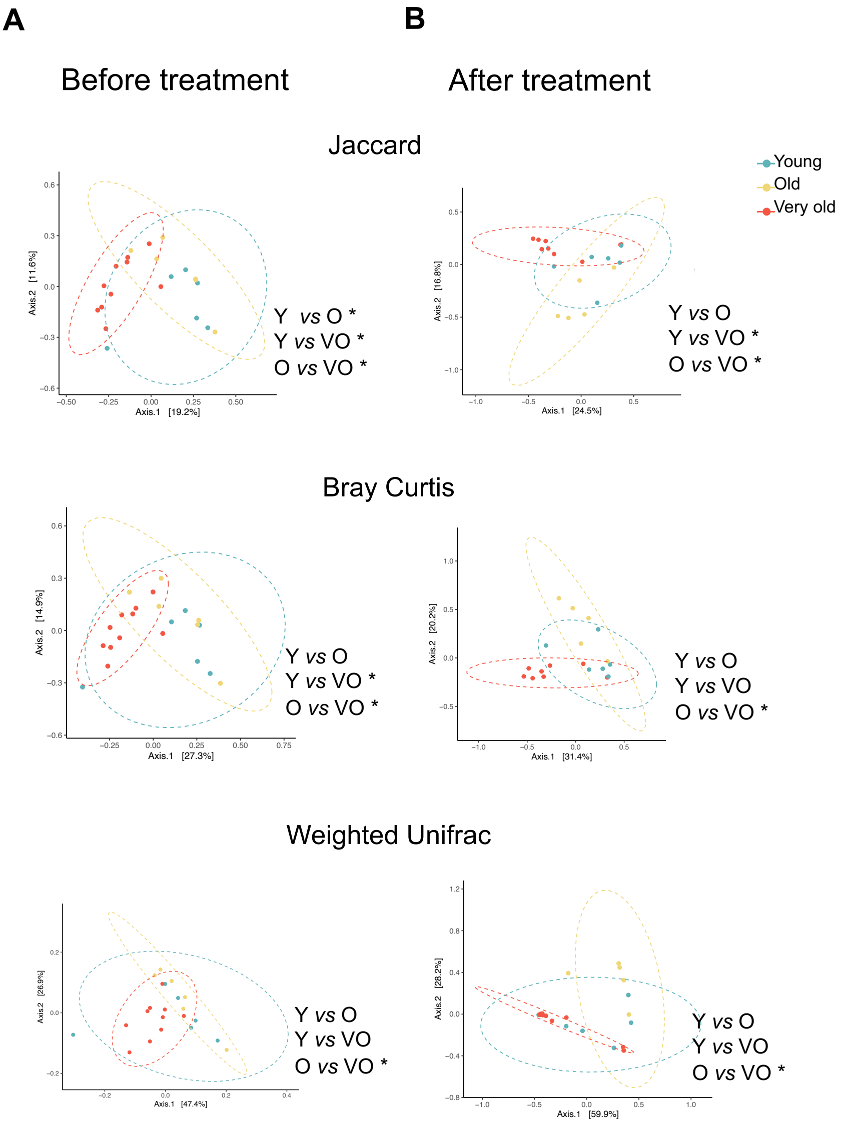

**Figure S1. Changes in Gut Microbiota Composition Before and After Treatment With Streptomycin and *E. coli* Colonization. Related to Figure 1.**
(A) and (B) NMDS illustrating beta-diversity (Jaccard, Bray Curtis, Weighted Unifrac) for young (n = 6) [S1], old (n = 6) [S1], and very old (n = 10) mice before (A) and after (B) treatment. Statistical comparisons were performed with PERMANOVA followed by pairwise post-hoc tests. *$P < 0.05$, ** $P < 0.01$, *** $P < 0.001$, **** $P < 0.0001$.

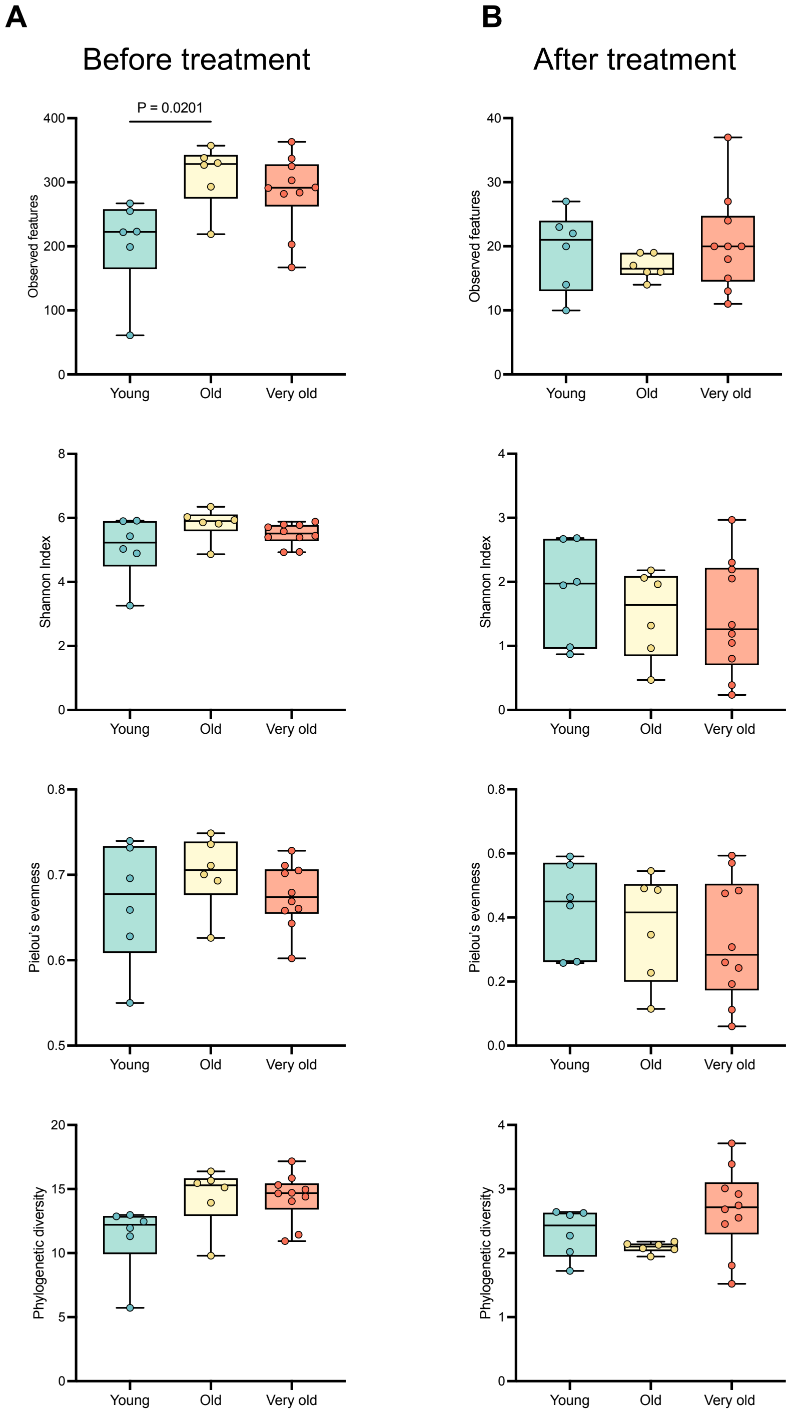

**Figure S2. Alpha Diversity Before and After Treatment with Streptomycin and *E. coli* Colonization. Related to Figure 1.**

(A) And (B) Alpha diversity measures (Observed features, Shannon Index, Pielou's evenness, Faith's phylogenetic diversity) of the gut microbiota of young (n = 6) [S1], old (n = 6) [S1], and very old (n = 10) mice before (A) and after (B) treatment. Differences in the median were tested using Kruskal Wallis test followed by Dunn's multiple comparisons correction.

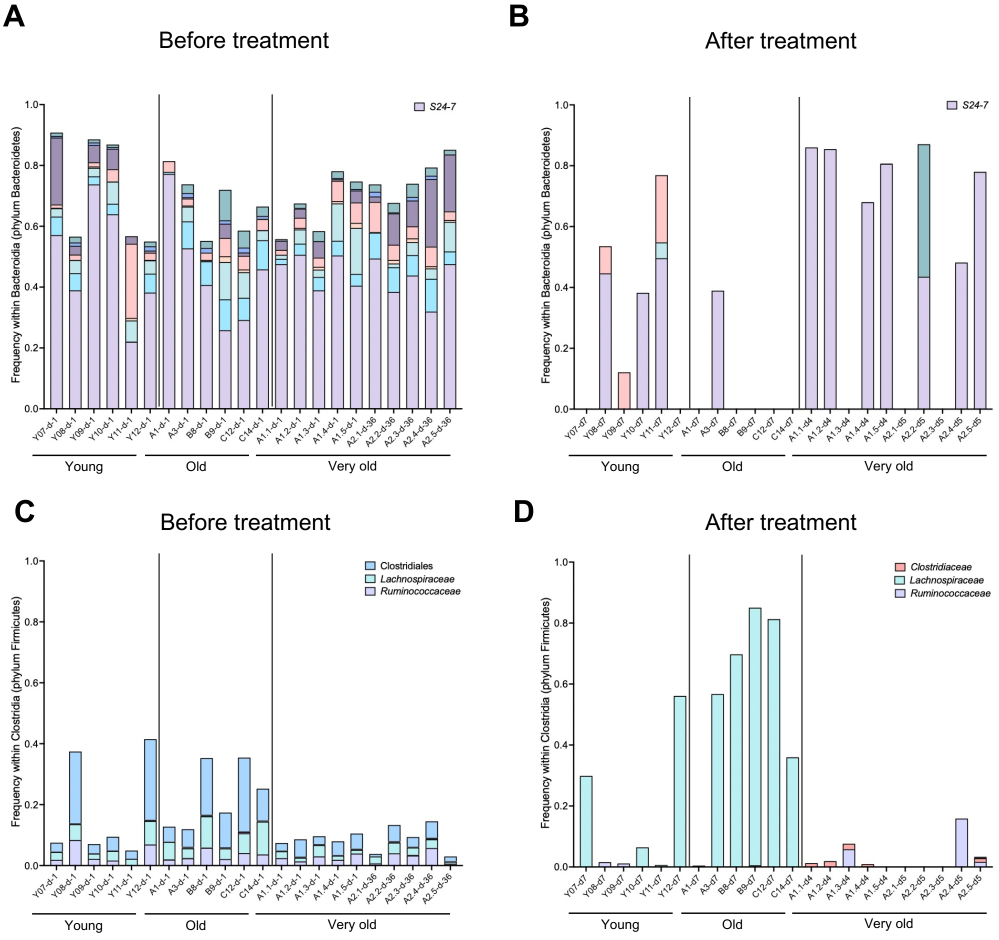

**Figure S3. Relative Frequency of Families Before and After Treatment with Streptomycin and *E. coli* Colonization. Related to Figure 1.**

(A) and (B) Taxa bar plots showing the relative frequency of bacterial families within the class Bacteroidia (Bacteroidetes phylum) before (A) and after (B) treatment in the gut of young (n = 6) [S1], old (n = 6) [S1], and very old (n = 10) mice.

(C) and (D) Taxa bar plots showing the relative frequency of bacterial families within the class Clostridia (Firmicutes phylum) before (C) and after (D) treatment in the gut of young (n = 6) [S1], old (n = 6) [S1], and very old (n = 10) mice.

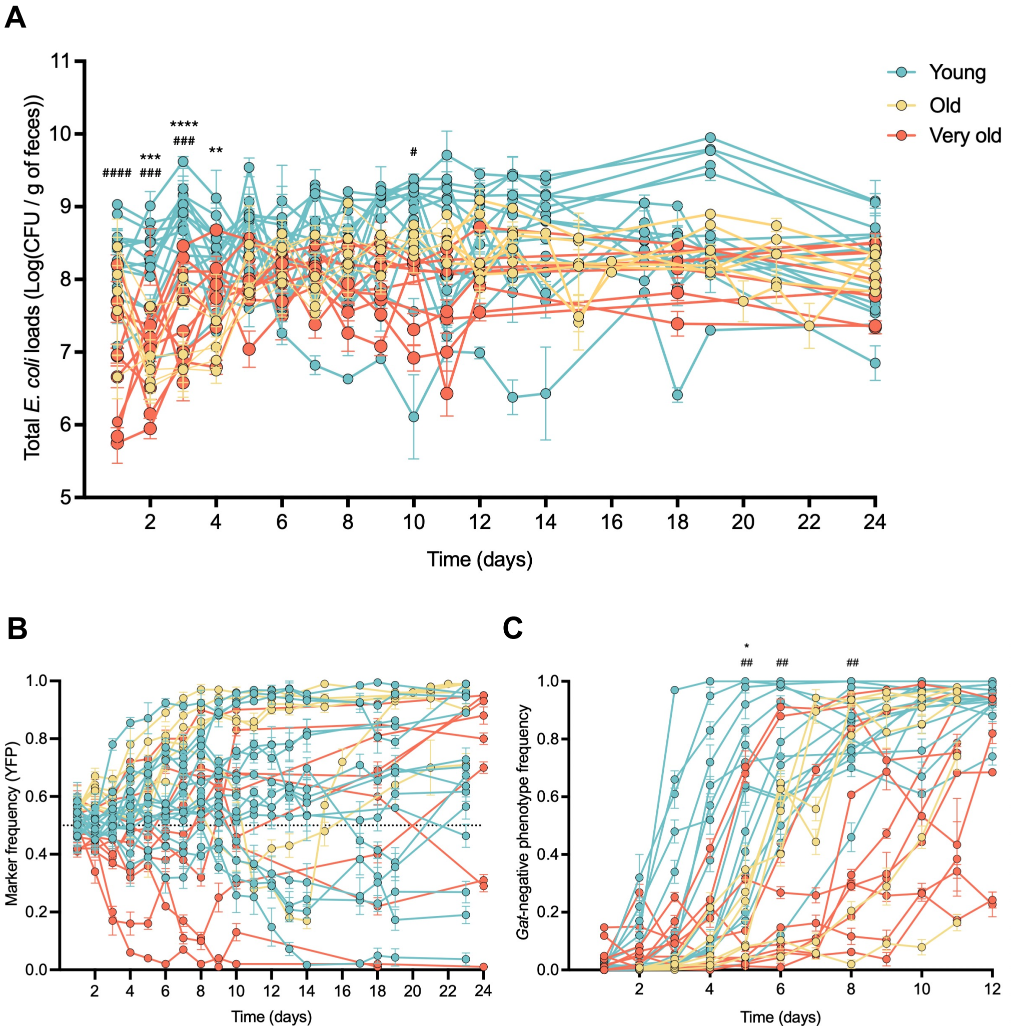

**Figure S4. Comparison of the Size of *E. Coli*'s Niche, the Pace of Evolution and the Emergence of a Highly Beneficial Phenotype (*Gat*-Negative). Related to Figure 2.**

(A) Total *E. coli* loads (per gram of feces) throughout the 24 days of adaptation of *E. coli* to the gut of young (n = 15) [S2], old (n = 6) [S1], and very old (n = 10) mice. Error bars represent the ± 2*SEM. The effects of the age and time on the loads were tested by a linear mixed model followed by a type III ANOVA and post-hoc contrasts with Bonferroni's correction. * — comparisons between young and old; # — comparisons between young and very old mice.

(B) Temporal dynamics of the frequency of the YFP-expressing *E. coli* populations evolving in young (n = 15) [S2], old (n = 6) [S1], and very old mice (n = 10).

(C) Emergence of the adaptive phenotype gat-negative in *E. coli* populations evolving in young (n = 15) [S2], old (n = 6) [S1], and very old mice (n = 10). The effects of the age and time on the *gat*-negative frequency were tested by a linear mixed model followed by a type III ANOVA and post-hoc contrasts with Bonferroni's correction. "*" refers to comparisons between young and old and "#" refers to comparisons between young and very old mice.

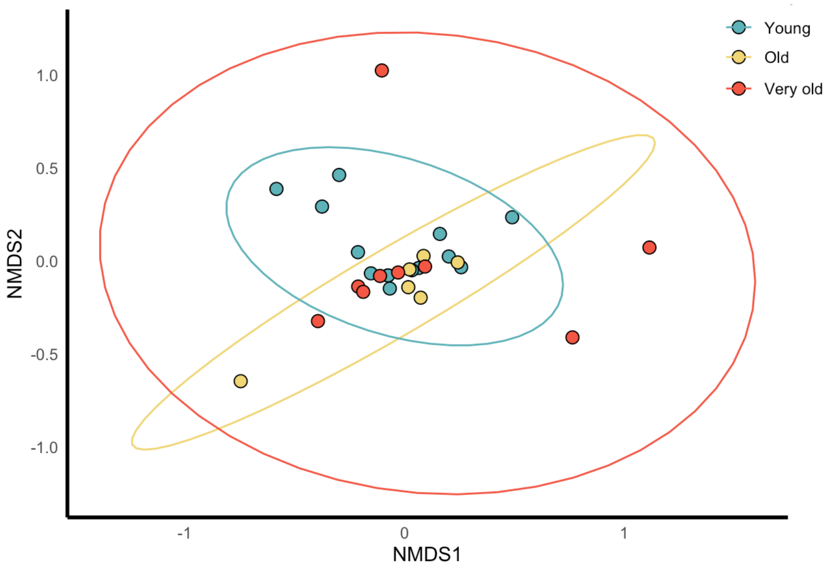

**Figure S5. Comparison of *E. coli's* Genetic Diversity Across Age Groups During Evolution. Related to Figure 2.**
NMDS analysis of gene diversity (Bray Curtis) between young (n = 14) [S1,S3], old (n = 6) [S1], and very old (n = 9) mice (PERMANOVA, *P* = 0.352).

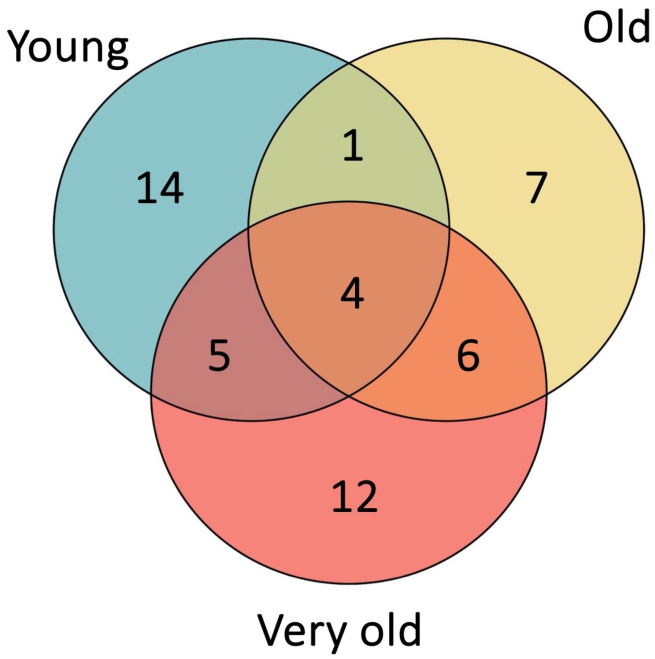

**Figure S6. Shared and Unique Gene Targets in *E. coli* Populations Across Age Groups at Day 11 and 24 of Evolution. Related to Figure 2.**
Venn diagram displaying overlap of the mutational targets (above a frequency of 5%) between young (n = 14) [S1,S3], old (n = 6) [S1], and very old (n = 9). The Venn diagram was produced using the R package *VennDiagram* [S4].

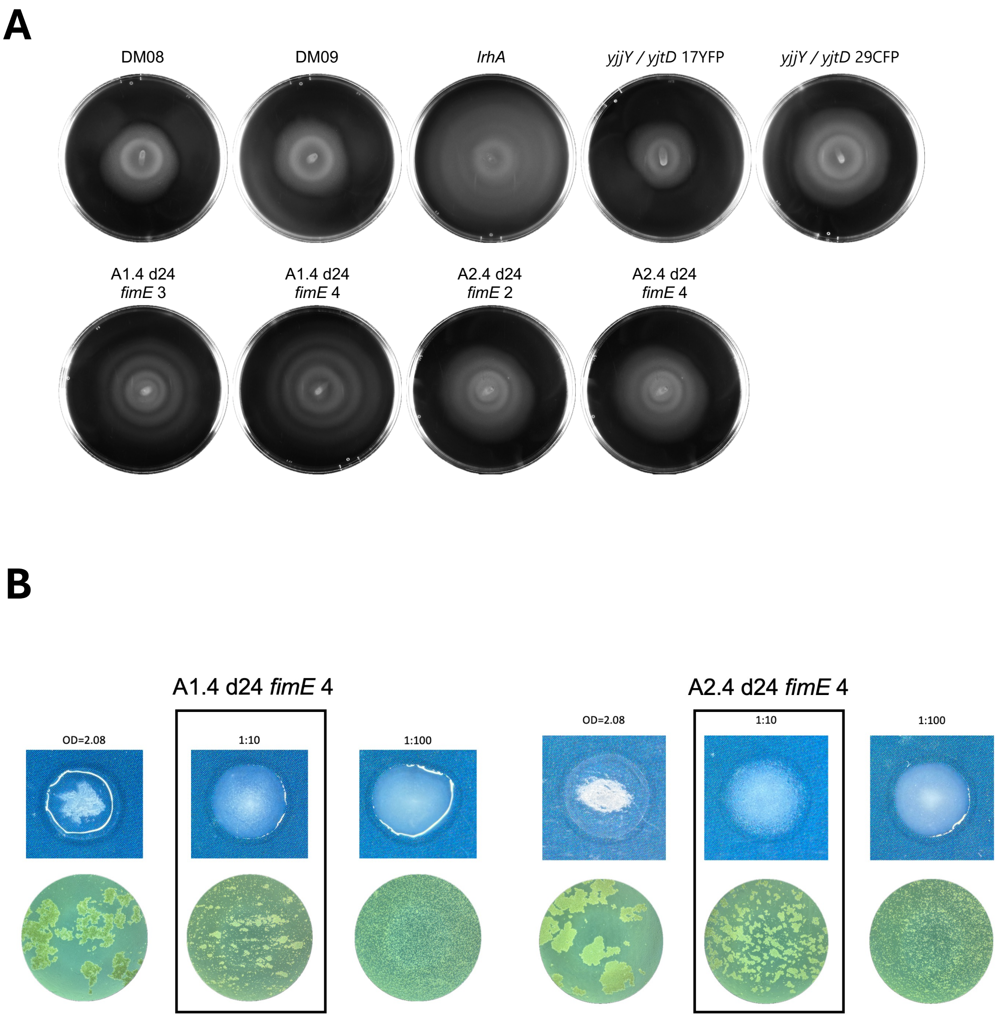

**Figure S7. The Aged Gut Selects for Hypermotility and Increased Adhesion. Related to Figure 3.**

(A) Representative images from the swarming motility of the ancestral (DM08 and DM09), *lrhA*, *yjjY*/*yjtD* and *fimE* clones. Swarm expansion, resulting from bacterial growth, appears in white, whereas uncolonized agar appears in black.

(B) Representative images (top, in blue: macroscopic image; bottom, in green: microscopic image) from the aggregates formed by interaction of E. coli with yeast, for the remaining two out of four *fimE* clones. This assay was performed in 2 independent replicates.

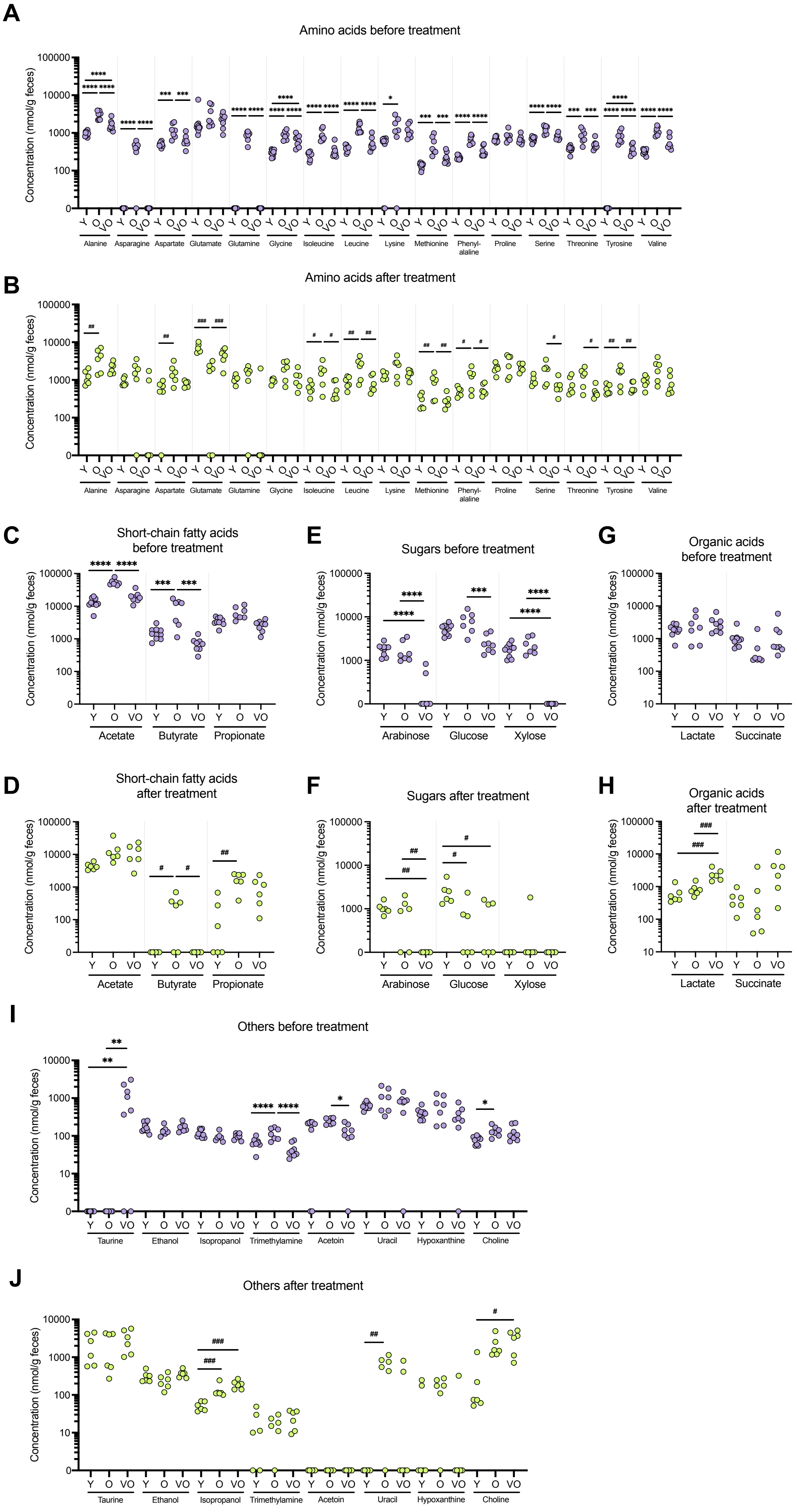
A
Amino acids before treatment
Concentration (nmol/g feces)
Y O VO
Alanine
Asparagine
Aspartate
Glutamate
Glutamine
Glycine
Isoleucine
Leucine
Lysine
Methionine
Phenyl-alaline
Proline
Serine
Threonine
Tyrosine
Valine
B
Amino acids after treatment
C
Short-chain fatty acids before treatment
Acetate
Butyrate
Propionate
D
Short-chain fatty acids after treatment
E
Sugars before treatment
Arabinose
Glucose
Xylose
F
Sugars after treatment
G
Organic acids before treatment
Lactate
Succinate
H
Organic acids after treatment
I
Others before treatment
Taurine
Ethanol
Isopropanol
Trimethylamine
Acetoin
Uracil
Hypoxanthine
Choline
J
Others after treatment

**Figure S8. Detailed Metabolite Levels in the Gut of Young, Old, and Very Old Mice Before and After Streptomycin Treatment and Colonization. Related to Figure 4.**
Scatter plots showing the concentrations of various metabolites before (purple circles) and after treatment (green circles) across the different age groups. (A,B) Amino acids; (C,D) Short-chain fatty acids; (E,F) Sugars; (G,H) Organic acids; (I,J) Other metabolites. Statistical comparisons were made on log-transformed values with One-way ANOVA followed by Tukey's multiple comparisons test. * – comparisons among age groups before treatment; # — comparisons among age groups after treatment. *$P$ < 0.05, **$P$ < 0.01, ***$P$ < 0.001, ****$P$ < 0.0001.

**Table S1. Differentially Abundant ASVs Before and After Streptomycin Treatment and Colonization. Related to Figure 1.** Differentially abundant ASVs in the gut of young (n = 6) [S1], old (n= 6) [S1], and very old (n = 10) mice before and after streptomycin treatment and *E. coli* colonization obtained from the analysis with ANCOM (threshold 0.8) [S5].

| | ASV | Taxonomical classification | Y *vs.* O | Y *vs.* VO | O *vs.* VO |
|---|---|---|---|---|---|
| Before treatment | 8f98fb8693ed59c21399d83ce2d10724 | *Akkermansia muciniphila* | | ↑ in VO (W=610) | ↑ in VO (W=630) |
| | b4d2dc7e79e032c6486253e7a49f074b | *Bacteroides acidifaciens* | ↑ in O (W=520) | ↑ in VO (W=553) | |
| | 583d040da1f83aa5e2797c3aa83a93b5 | Clostridiales (1) | | | ↑ in VO (W=505) |
| | b2bb76e45be2318c640042739a205314 | Clostridiales (2) | | | ↑ in VO (W=530) |
| | 6e61f4bede587c68efcaa016e92085f2 | Clostridiales (3) | | | ↑ in O (W=626) |
| | 36934aef9f30b3a823b34b6d0e6b6d89 | Clostridiales (4) | | | ↑ in O (W=616) |
| | cda4d243ad8b0e0814bd96abfbc9f6d5 | Clostridiales (5) | | | ↑ in O (W=537) |
| | 18d3f11b4f13f904a2dde91f7147c2dd | Clostridiales (6) | | | ↑ in O (W=529) |
| | 41e7e5b9b1734e7c70e403b282379dac | *Coprococcus* | | | ↑ in VO (W=545) |
| | 7b922bd11ea3f2de8a1d01bc0314871f | *Neisseria* | | | ↑ in VO (W=545) |
| | ea058e1daf36b416ff33f8590ced424c | *Oscillospira* | | ↑ in VO (W=561) | ↑ in VO (W=615) |
| | 901e24ec7d14fc177f7f9c15ab0f972d | *Prevotella* | ↑ in Y (W=561) | ↑ in Y (W=559) | |
| | 0b98a395ba83623146746156a838174e | RF32 | | | ↑ in VO (W=546) |
| | cfd65c7bbc3926cdc9b88c2e38bbecaa | *Rikenellaceae* | ↑ in Y (W=497) | | ↑ in VO (W=628) |
| | b61ee4ecb0582e6350d1148cb523de88 | *S24-7* (1) | ↑ in O (W=479) | | |
| | 2db69eb5d23a7acdbe62c107a23acf2b | *S24-7* (2) | | ↑ in VO (W=562) | ↑ in VO (W=605) |
| | 47a33be537ddd236047d0762a0289582 | *S24-7* (3) | | ↑ in VO (W=557) | |
| | c1a01df6711439c8991237b3b9838055 | *S24-7* (4) | | ↑ in VO (W=533) | ↑ in VO (W=583) |
| | 5b2af6c151f46e308f995abe9e3eb50e | *S24-7* (5) | | | ↑ in VO (W=623) |
| | d23fbef2f31d48eda40876cdbc49933a | *Staphylococcus* | | ↑ in Y (W=506) | |
| | 52fdc469d22151ff5173d5f656fbbf43 | *Sutterella* (1) | ↑ in Y (W=515) | | ↑ in VO (W=641) |
| | 7d17d197d7978be0b6684209ce257e20 | *Sutterella* (2) | | ↑ in VO (W=594) | |
| | d353248c8a7d31bc9f8378a320b77924 | *Blautia producta* | | | ↑ in O (W=57) |
| | f0607d2c57afe6bb25bd64afa9869e28 | *Clostridiaceae* | | | ↑ in O (W=46) |

| | | | | | |
|---|---|---|---|---|---|
| | b85eba4bd049b34b9ab06853403f6675 | *Clostridium* (1) | | | ↑ in O (W=51) |
| | 9baebca868145d4bb01f9a4d22a434b3 | *Clostridium* (2) | | | ↑ in O (W=50) |
| | 22c8e6e55e7d3f64fc2fdf7b5cc181a2 | *Clostridium aldenense* | | | ↑ in O (W=52) |
| | 7b922bd11ea3f2de8a1d01bc0314871f | *Neisseria* | | ↑ in VO (W=51) | ↑ in VO (W=49) |
| After treatment | 38bfbd3a2cf83da8fe03cea0f6b4f5f3 | *Pseudomonas fragi* | | | ↑ in O (W=46) |
| | 3b482b3f9da81db80a221a66031aa159 | *S24-7* (6) | | ↑ in VO (W=59) | ↑ in VO (W=57) |
| | ff4ae0f811e8dcb2d76d7d8acc73deb8 | *S24-7* (7) | | ↑ in VO (W=52) | ↑ in VO (W=52) |
| | 686b1c1752a35be01421dfdacfa244e1 | *S24-7* (8) | | | ↑ in VO (W=56) |
| | d23fbef2f31d48eda40876cdbc49933a | *Staphylococcus* | ↑ in Y (W=51) | | ↑ in O (W=59) |
| | 5608c3e6c9de9ceb79610e7786bd0ac4 | *Veilonella dispar* | | ↑ in VO (W=47) | ↑ in VO (W=46) |

**Table S2. Mutations Observed in *E. coli* Populations After 11 Days of Evolution in Very Old Mice. Related to Figures 2 and 3.** Adaptive mutations observed in independently evolved populations of *E. coli* after 11 days in very old mice. SNPs are represented by an arrow between the ancestral and the evolved nucleotide. Δ – deletion event; + – insertion of nucleotides; IS – insertion sequence at the indicated position. 1→2 – duplication.

| Population | Genome position | Gene | Mutation | Annotation | Frequency |
|---|---|---|---|---|---|
| A1.1 | 405692 | *aroL* → | C→A | L22I (CTT→ATT) | 3% |
| | 415553 | *sbcD* ← | C→T | P208P (CCG→CCA) | 2% |
| | 863839 | *moeB* ← | A→G | C172R (TGC→CGC) | 10% |
| | 1044369 | *gfcE* ← | C→T | V220I (GTA→ATA) | 3% |
| | 1067723 | *ymdF* → / ← *rutG* | C→T | intergenic (+246/+11) | 5% |
| | 1195443 | *e14 prophage* | Δ15,192 bp | e14 prophage loss | 4% |
| | 1296579 | *adhE* ← | Δ3 bp | coding (764-766/2676 nt) | 1% |
| | 1535191 | *narW* ← | $(AGC)_{2\rightarrow 1}$ | coding (141-143/696 nt) | 2% |
| | 1720637 | *ydhK* → | Δ3 bp | coding (493-495/2013 nt) | 3% |
| | 1854063 | *ydjH* ← | G→T | T297N (ACC→AAC) | 4% |
| | 2173891 | *gatZ* ← | Δ5 bp | coding (449-453/1263 nt) | 39% |
| | 2175204 | *gatY* ← | IS*1* (+) +9 bp | coding (15-23/855 nt) | 7% |
| | 2634607 | *der* ← | C→T | A258T (GCT→ACT) | 2% |
| | 2718071 | *pka* → | C→T | R33C (CGT→TGT) | 4% |

| | | | | | |
|---|---|---|---|---|---|
| | 3756090 | *selB* ← | IS*1* (–) +9 bp | coding (1787-1795/1845 nt) | 7% |
| | 3756253 | *selB* ← | +TAACGATCTT | coding (1632/1845 nt) | 41% |
| | 4082473 | *fdoG* ← | C→T | G458E (GGG→GAG) | 55% |
| **A1.2** | 1195443 | *e14 prophage* | Δ15,192 bp | e14 prophage loss | 7% |
| | 1737784 | *ydhB* ← | T→C | V13V (GTA→GTG) | 2% |
| | 2175295 | *gatY* ← / ← *fbaB* | IS*1* (–) +9 bp | intergenic (-69/+231) | 12% |
| | 3123436 | *glcF* ← | Δ2 bp | coding (46/1224 nt) | 4% |
| | 3968441 | *wecB* → | Δ3 bp | coding (286-288/1131 nt) | 1% |
| | 4592985 | *yjjM* ← | IS*5* (–) +4 bp | coding (887-890/915 nt) | 2% |
| | 4638639 | *yjjY* → / → *yjtD* | IS*2* (–) +5 bp | intergenic (+74/-322) | 5% |
| | 4638670 | *yjjY* → / → *yjtD* | IS*5* (–) +4 bp | intergenic (+105/-292) | 28% |
| | 4638670 | *yjjY* → / → *yjtD* | IS*5* (+) +4 bp | intergenic (+105/-292) | 11% |
| | 4638719 | *yjjY* → / → *yjtD* | IS*5* (–) +4 bp | intergenic (+154/-243) | 10% |
| | 4638762 | *yjjY* → / → *yjtD* | IS*1* (+) +9 bp | intergenic (+197/-195) | 21% |
| | 4638771 | *yjjY* → / → *yjtD* | IS*5* (+) +4 bp | intergenic (+206/-191) | 19% |
| **A1.3** | 1195443 | *e14 prophage* | Δ15,192 bp | e14 prophage loss | 3% |
| | 1720637 | *ydhK* → | Δ3 bp | coding (493-495/2013 nt) | 2% |
| | 2172673 | *gatA* ← | IS*1* (+) +8 bp | coding (392-399/453 nt) | 11% |
| | 2173249 | *gatZ* ← | IS*1* (+) +9 bp | coding (1087-1095/1263 nt) | 19% |
| | 2173306 | *gatZ* ← | IS*1* (+) +9 bp | coding (1030-1038/1263 nt) | 75% |
| | 2173905 | *gatZ* ← | Δ5 bp | coding (435-439/1263 nt) | 2% |
| | 2763434 | *yfjL* ← / ← *yfjM* | Δ8 bp | intergenic (-259/+94) | 2% |
| | 3465823 | *chiA* ← | C→T | D685N (GAC→AAC) | 2% |
| | 3756090 | *selB* ← | IS*1* (–) +9 bp | coding (1787-1795/1845 nt) | 6% |
| | 3756090 | *selB* ← | IS*1* (+) +9 bp | coding (1787-1795/1845 nt) | 18% |
| | 3758884 | *selA* ← | C→A | R130L (CGC→CTC) | 63% |
| | 3758913 | *selA* ← | A→C | Y120* (TAT→TAG) | 4% |

| | | | | | |
|---|---|---|---|---|---|
| | 4178685 | *rplL* → | Δ3 bp | coding (103-105/366 nt) | 1% |
| | 4430075 | *ytfE* ← / ← *ytfF* | C→A | intergenic (-69/+39) | 3% |
| | 1195443 | *e14 prophage* | Δ15,192 bp | e14 prophage loss | 4% |
| | 1291986 | *hns* ← | G→T | R54S (CGC→AGC) | 25% |
| | 1292064 | *hns* ← | (TTCCAGCATTTC)$_{1\rightarrow 2}$ | coding (82/414 nt) | 1% |
| | 1718955 | *slyA* ← / → *ydhI* | G→A | intergenic (-107/-94) | 14% |
| | 1835828 | *ynjC* → | T→A | V198D (GTT→GAT) | 4% |
| | 2172866 | *gatA* ← | IS*5* (–) +4 bp | coding (203-206/453 nt) | 5% |
| | 2175295 | *gatY* ← / ← *fbaB* | IS*1* (–) +9 bp | intergenic (-69/+231) | 7% |
| | 2659862 | *iscR* ← | A→T | C98S (TGC→AGC) | 8% |
| | 2827134 | *srlR* → | +GAAGA | coding (66/774 nt) | 2% |
| | 3123436 | *glcF* ← | Δ2 bp | coding (46/1224 nt) | 5% |
| | 3688311 | *bcsB* ← | Δ3 bp | coding (2318-2320/2340 nt) | 1% |
| | 4638628 | *yjjY* → / → *yjtD* | IS*2* (–) +5 bp | intergenic (+63/-333) | 6% |
| | 4638637 | *yjjY* → / → *yjtD* | IS*5* (+) +4 bp | intergenic (+72/-325) | 5% |
| | 4638670 | *yjjY* → / → *yjtD* | IS*5* (–) +4 bp | intergenic (+105/-292) | 6% |
| | 4638753 | *yjjY* → / → *yjtD* | IS*2* (–) +5 bp | intergenic (+188/-208) | 1% |
| **A1.4** | 4638771 | *yjjY* → / → *yjtD* | IS*5* (+) +4 bp | intergenic (+206/-191) | 10% |
| | 458014 | *clpX* → / → *lon* | IS*186* (–) +6 bp :: Δ1 bp | intergenic (+90/-93) | 1% |
| | 1195443 | *e14 prophage* | Δ15,192 bp | e14 prophage loss | 3% |
| | 1720637 | *ydhK* → | Δ3 bp | coding (493-495/2013 nt) | 2% |
| | 1727550 | *lhr* → | C→A | P147Q (CCG→CAG) | 4% |
| **A1.5** | 1824654 | *astE* ← | G→A | C98C (TGC→TGT) | 4% |
| | 2175295 | *gatY* ← / ← *fbaB* | IS*1* (–) +9 bp | intergenic (-69/+231) | 66% |
| | 3091759 | *yqgF* → | G→A | A80T (GCC→ACC) | 3% |
| | 3734676 | *bax* ← | Δ3 bp | coding (523-525/825 nt) | 1% |
| | 4166219 | *rrsB* → | C→A | noncoding (1538/1542 nt) | 7% |

| | | | | | |
|---|---|---|---|---|---|
| A2.1 | 382402 | *yaiP* ← | Δ1 bp | coding (758/1197 nt) | 2% |
| | 759145 | *sucA* → | +GTACT | coding (1217/2802 nt) | 16% |
| | 819824 | *ybhL* → / → *ybhM* | Δ1 bp | intergenic (+13/-192) | 2% |
| | 1195443 | *e14 prophage* | Δ15,192 bp | e14 prophage loss | 9% |
| | 1264657 | *prfA* → | C→A | A141A (GCC→GCA) | 2% |
| | 1610599 | *glsB* ← | +CC | coding (677/927 nt) | 3% |
| | 2172866 | *gatA* ← | IS*5* (–) +4 bp | coding (203-206/453 nt) | 6% |
| | 2173261 | *gatZ* ← | IS*1* (+) +8 bp | coding (1076-1083/1263 nt) | 6% |
| | 2173546 | *gatZ* ← | IS*1* (+) +9 bp | coding (790-798/1263 nt) | 2% |
| | 2173900 | *gatZ* ← | A→C | L148L (CTT→CTG) | 3% |
| | 2173905 | *gatZ* ← | Δ5 bp | coding (435-439/1263 nt) | 24% |
| | 2175233 | *gatY* ← / ← *fbaB* | IS*2* (+) +5 bp | intergenic (-7/+297) | 25% |
| | 2720768 | *pssA* → | G→A | R7H (CGT→CAT) | 2% |
| | 2937638 | *fucR* → | Δ2 bp | coding (249/732 nt) | 2% |
| | 3123436 | *glcF* ← | Δ2 bp | coding (46/1224 nt) | 5% |
| | 3174366 | *cpdA* ← | Δ3 bp | coding (488-490/828 nt) | 1% |
| | 3652075 | *slp* → | $(AACC)_{2→1}$ | coding (92-95/567 nt) | 0% |
| | 3688311 | *bcsB* ← | Δ3 bp | coding (2318-2320/2340 nt) | 1% |
| | 3734676 | *bax* ← | Δ3 bp | coding (523-525/825 nt) | 2% |
| | 4638771 | *yjjY* → / → *yjtD* | IS*5* (–) +4 bp | intergenic (+206/-191) | 2% |
| | 4638842:1 | *yjjY* → / → *yjtD* | +T | intergenic (+277/-123) | 94% |
| A2.2 | 293096 | *yagL* ← | IS*2* (–) +5 bp | coding (43-47/699 nt) | 2% |
| | 1090815 | *pgaA* ← | G→A | A233V (GCC→GTC) | 1% |
| | 1195443 | *e14 prophage* | Δ15,192 bp | e14 prophage loss | 3% |
| | 1292052 | *hns* ← | +TTCCAGCATTTC | coding (94/414 nt) | 9% |
| | 1907822 | *kdgR* ← | G→A | S101F (TCC→TTC) | 2% |
| | 2172033 | *gatC* ← | Δ5 bp | coding (264-268/1356 nt) | 1% |

| | | | | | |
|---|---|---|---|---|---|
| | 2173905 | *gatZ* ← | Δ5 bp | coding (435-439/1263 nt) | 2% |
| | 2175299 | *gatY* ← / ← *fbaB* | IS*2* (+) +5 bp | intergenic (-73/+231) | 21% |
| | 3565643 | *glgA* ← | C→T | W138* (TGG→TGA) | 2% |
| | 4043118 | *yihF* → | Δ20 bp | coding (897-916/1431 nt) | 1% |
| | 4638628 | *yjjY* → / → *yjtD* | IS*2* (–) +5 bp | intergenic (+63/-333) | 57% |
| | 4638670 | *yjjY* → / → *yjtD* | IS*5* (–) +4 bp | intergenic (+105/-292) | 19% |
| | 4638771 | *yjjY* → / → *yjtD* | IS*5* (+) +4 bp | intergenic (+206/-191) | 20% |
| **A2.3** | 1195443 | *e14 prophage* | Δ15,192 bp | e14 prophage loss | 9% |
| | 1296579 | *adhE* ← | Δ3 bp | coding (764-766/2676 nt) | 2% |
| | 1444920 | *feaR* ← | A→T | F130I (TTT→ATT) | 4% |
| | 1741614 | *mdtK* → | C→T | A45V (GCG→GTG) | 2% |
| | 1959830 | *argS* → / → *yecT* | G→A | intergenic (+11/-166) | 2% |
| | 2173891 | *gatZ* ← | Δ5 bp | coding (449-453/1263 nt) | 12% |
| | 2174867 | *gatY* ← | IS*4* (+) +12 bp | coding (349-360/855 nt) | 6% |
| | 2175256 | *gatY* ← / ← *fbaB* | C→A | intergenic (-30/+278) | 17% |
| | 2175267 | *gatY* ← / ← *fbaB* | G→T | intergenic (-41/+267) | 14% |
| | 3123436 | *glcF* ← | Δ2 bp | coding (46/1224 nt) | 4% |
| | 3529475 | *yhgE* ← | C→T | V329V (GTG→GTA) | 3% |
| | 3649418 | *yhiS* → | G→A | pseudogene (105/741 nt) | 1% |
| | 4178685 | *rplL* → | Δ3 bp | coding (103-105/366 nt) | 1% |
| | 4298875 | *mdtP* ← | $(CAC)_{2\rightarrow1}$ | coding (177-179/1467 nt) | 2% |
| | 4638628 | *yjjY* → / → *yjtD* | IS*2* (–) +5 bp | intergenic (+63/-333) | 56% |
| | 4638637 | *yjjY* → / → *yjtD* | IS*5* (+) +4 bp | intergenic (+72/-325) | 83% |
| | 4638670 | *yjjY* → / → *yjtD* | IS*5* (–) +4 bp | intergenic (+105/-292) | 3% |
| | 4638771 | *yjjY* → / → *yjtD* | IS*5* (+) +4 bp | intergenic (+206/-191) | 5% |
| **A2.4** | 784026 | *zitB* ← | Δ6 bp | coding (16-21/942 nt) | 1% |
| | 1195443 | *e14 prophage* | Δ15,192 bp | e14 prophage loss | 3% |

| | | | | | |
|---|---|---|---|---|---|
| | 1720637 | *ydhK* → | Δ3 bp | coding (493-495/2013 nt) | 3% |
| | 1907380 | *kdgR* ← | +GC | coding (744/792 nt) | 5% |
| | 1983600 | *araF* ← | Δ1 bp | coding (553/990 nt) | 13% |
| | 2172334 | *gatB* ← | IS*1* (–) +9 bp | coding (247-255/285 nt) | 14% |
| | 2172338 | *gatB* ← | IS*1* (+) +9 bp | coding (243-251/285 nt) | 8% |
| | 2172662 | *gatA* ← | IS*1* (+) +9 bp | coding (402-410/453 nt) | 15% |
| | 2172684 | *gatA* ← | IS*5* (–) +4 bp | coding (385-388/453 nt) | 18% |
| | 2172755 | *gatA* ← | IS*1* (–) +8 bp | coding (310-317/453 nt) | 9% |
| | 2172882 | *gatA* ← | IS*1* (+) +9 bp | coding (182-190/453 nt) | 6% |
| | 2173327 | *gatZ* ← | IS*1* (–) +8 bp | coding (1010-1017/1263 nt) | 1% |
| | 2173471 | *gatZ* ← | IS*1* (–) +9 bp | coding (865-873/1263 nt) | 5% |
| | 2173522 | *gatZ* ← | IS*1* (+) +9 bp | coding (814-822/1263 nt) | 9% |
| | 2173891 | *gatZ* ← | Δ5 bp | coding (449-453/1263 nt) | 1% |
| | 2175283 | *gatY* ← / ← *fbaB* | IS*1* (–) +9 bp | intergenic (-57/+243) | 6% |
| | 2624837 | *yfgF* ← | T→C | E708E (GAA→GAG) | 1% |
| | 2771809 | *yfjW* → | T→C | I157T (ATA→ACA) | 1% |
| | 2827555 | *srlR* → | IS*3* (–) +5 bp :: Δ3 bp +T | coding (487-491/774 nt) | 3% |
| | 3252152 | *yhaJ* ← | Δ3 bp | coding (83-85/897 nt) | 1% |
| | 3785915 | *envC* → | Δ3 bp | coding (1055-1057/1260 nt) | 2% |
| | 3999676 | *corA* → | Δ3 bp | coding (228-230/951 nt) | 1% |
| | 4354001 | *yjdL* ← | G→A | A145V (GCG→GTG) | 2% |
| | 4512268 | *fecB* ← | C→T | A22T (GCC→ACC) | 2% |
| | 4593214 | *yjjM* ← | G→A | L221F (CTT→TTT) | 2% |
| | 4593422 | *yjjM* ← | T→A | Q151H (CAA→CAT) | 3% |
| **A2.5** | 203011 | *lpxA* → | C→T | A151V (GCG→GTG) | 2% |
| | 223673 | *gmhB* → / → *rrsH* | A→G | intergenic (+265/-98) | 15% |
| | 508154 | *copA* ← | G→A | S817L (TCG→TTG) | 2% |

| | | | | | |
|---|---|---|---|---|---|
| | 953896 | *focA* ← / ← *ycaO* | IS*5* (+) Δ5 bp | intergenic (-207/+195) | 1% |
| | 1195443 | *e14 prophage* | Δ15,192 bp | e14 prophage loss | 15% |
| | 1198930 | *intE* ← | C→G | G367A (GGT→GCT) | 35% |
| | 2172214 | *gatC* ← | IS*1* (+) +8 bp | coding (80-87/1356 nt) | 4% |
| | 2172334 | *gatB* ← | IS*1* (–) +9 bp | coding (247-255/285 nt) | 2% |
| | 2172755 | *gatA* ← | IS*1* (–) +8 bp | coding (310-317/453 nt) | 38% |
| | 2173522 | *gatZ* ← | IS*1* (–) +9 bp | coding (814-822/1263 nt) | 2% |
| | 2173993 | *gatZ* ← | IS*1* (+) +9 bp | coding (343-351/1263 nt) | 8% |
| | 2827297 | *srlR* → | A→T | K77* (AAG→TAG) | 28% |
| | 2827492 | *srlR* → | G→A | G142S (GGC→AGC) | 32% |
| | 3026201 | *ygfQ* → | T→C | I353I (ATT→ATC) | 2% |
| | 3453566 | *gspA* ← / → *gsp C* | +ACCC | intergenic (-146/-34) | 27% |
| | 3516346 | *aroB* ← | $(CGC)_{2\rightarrow 1}$ | coding (161-163/1089 nt) | 1% |
| | 4495767 | *intB* → | Δ3 bp | pseudogene (1070-1072/1266 n t) | 1% |
| | 4601134 | *yjjP* ← / → *yjjQ* | IS*5* (–) +4 bp | intergenic (-253/-363) | 1% |

**Table S3. Mutations Observed in *E. coli* Populations After 24 Days of Evolution in Very Old Mice. Related to Figures 2 and 3.** Adaptive mutations observed in independently evolved populations of *E. coli* after 24 days in very old mice. SNPs are represented by an arrow between the ancestral and the evolved nucleotide. Δ – deletion event; + – insertion of nucleotides; IS – insertion sequence at the indicated position. 1→2 – duplication.

| Population | Genome position | Gene | Mutation | Annotation | Frequency |
|---|---|---|---|---|---|
| A1.1 | 18284 | *nhaA* → | C→T | L266L (CTG→TTG) | 15% |
| | 722320 | *kdpD* ← | Δ3 bp | coding (1316-1318/2685 nt) | 3% |
| | 1195443 | *e14 prophage* | Δ15,192 bp | e14 prophage loss | 6% |
| | 137723 | *appC* → | C→T | A254V (GCG→GTG) | 4% |
| | 2173522 | *gatZ* ← | IS*1* (–) +9 bp | coding (814-822/1263 nt) | 3% |
| | 2173891 | *gatZ* ← | Δ5 bp | coding (449-453/1263 nt) | 15% |
| | 2174058 | *gatZ* ← | G→A | Q96* (CAA→TAA) | 8% |
| | 2174503 | *gatY* ← | IS*4* (+) +11 bp | coding (714-724/855 nt) | 5% |
| | 2175204 | *gatY* ← | IS*1* (+) +9 bp | coding (15-23/855 nt) | 8% |
| | 2701640 | *rnc* ← | G→T | P149Q (CCA→CAA) | 5% |
| | 3756253 | *selB* ← | +TAACGATCTT | coding (1632/1845 nt) | 62% |
| | 4082473 | *fdoG* ← | C→T | G458E (GGG→GAG) | 34% |
| | 4638637 | *yjjY* → / → *yjtD* | IS*5* (+) +4 bp | intergenic (+72/-325) | 4% |
| A1.2 | 231184 | *yafD* → | G→A | P21P (CCG→CCA) | 1% |
| | 582299 | *tfaX* → | C→T | pseudogene (124/183 nt) | 1% |
| | 1195443 | *e14 prophage* | Δ15,192 bp | e14 prophage loss | 7% |
| | 1736219 | *purR* → | G→T | G118C (GGT→TGT) | 3% |
| | 1988099 | *tyrP* → | G→A | G132D (GGT→GAT) | 2% |
| | 2172772 | *gatA* ← | IS*1* (–) +9 bp | coding (292-300/453 nt) | 7% |
| | 2175243 | *gatY* ← / ← *fbaB* | T→G | intergenic (-17/+291) | 12% |
| | 2175295 | *gatY* ← / ← *fbaB* | IS*1* (–) +9 bp | intergenic (-69/+231) | 80% |
| | 2175295 | *gatY* ← / ← *fbaB* | Δ1 bp | intergenic (-69/+239) | 9% |
| | 2175296 | *gatY* ← / ← *fbaB* | Δ1 bp | intergenic (-70/+238) | 9% |
| | 2326978 | *yfaQ* ← | G→T | P281H (CCC→CAC) | 3% |
| | 2404461 | *lrhA* ← | G→A | T68I (ACT→ATT) | 6% |

| | | | | | |
|---|---|---|---|---|---|
| | 2404571 | *lrhA* ← | Δ9 bp | coding (85-93/939 nt) | 27% |
| | 2404571 | *lrhA* ← | (GCGGCAGCT)$_{1\rightarrow 2}$ | coding (93/939 nt) | 20% |
| | 2404817 | *lrhA* ← / → *alaA* | IS*1* (+) +9 bp | intergenic (-154/-758) | 2% |
| | 2888639 | *cysJ* ← | +GTCAGC | coding (1282/1800 nt) | 3% |
| | 2891037 | *ygcN* → | T→C | I120T (ATC→ACC) | 2% |
| | 4358139 | *cadB* ← / ← *cad C* | IS*2* (–) +5 bp | intergenic (-85/+276) | 1% |
| | 4592985 | *yjjM* ← | IS*5* (–) +4 bp | coding (887-890/915 nt) | 6% |
| | 4638762 | *yjjY* → / → *yjtD* | IS*1* (+) +9 bp | intergenic (+197/-195) | 1% |
| | 532268 | *allR* → | C→T | Q12* (CAG→TAG) | 15% |
| | 669769 | *nadD* ← | G→A | G9G (GGC→GGT) | 2% |
| | 1195443 | *e14 prophage* | Δ15,192 bp | e14 prophage loss | 5% |
| | 1292064 | *hns* ← | (TTCCAGCATTTC)$_{1\rightarrow 2}$ | coding (82/414 nt) | 46% |
| | 1692993 | *uidA* ← | C→T | G368D (GGC→GAC) | 3% |
| | 2172931 | *gatA* ← | IS*1* (–) +8 bp | coding (134-141/453 nt) | 2% |
| | 2173014 | *gatA* ← | IS*5* (–) +4 bp | coding (55-58/453 nt) | 2% |
| | 2173249 | *gatZ* ← | IS*1* (+) +9 bp | coding (1087-1095/1263 nt) | 7% |
| | 2173306 | *gatZ* ← | IS*1* (+) +9 bp | coding (1030-1038/1263 nt) | 19% |
| **A1.3** | 2173905 | *gatZ* ← | Δ5 bp | coding (435-439/1263 nt) | 10% |
| | 2814351 | *gshA* ← | G→A | G37G (GGC→GGT) | 2% |
| | 2819391 | *alaS* ← | C→T | V215I (GTC→ATC) | 3% |
| | 3265169 | *tdcA* ← / → *tdcR* | IS*5* (+) +4 bp | intergenic (-82/-230) | 39% |
| | 3756090 | *selB* ← | IS*1* (–) +9 bp | coding (1787-1795/1845 nt) | 8% |
| | 3757503 | *selB* ← | G→A | Q128* (CAG→TAG) | 47% |
| | 3758250 | *selA* ← | +T | coding (1023/1392 nt) | 14% |
| | 3758884 | *selA* ← | C→A | R130L (CGC→CTC) | 18% |
| | 3758913 | *selA* ← | A→C | Y120* (TAT→TAG) | 4% |

| | | | | | |
|---|---|---|---|---|---|
| | 4073139 | *yihW* → | (CCCGCGCTTTGAAGTGATGGTGCCCCGGCGGTACGTTGC)$_{1\rightarrow 2}$ | coding (448/786 nt) | 5% |
| | 4593009 | *yjjM* ← | IS*5* (–) +4 bp | coding (863-866/915 nt) | 1% |
| | 4593048 | *yjjM* ← | T→C | D276G (GAT→GGT) | 14% |
| | 4638628 | *yjjY* → / → *yjtD* | IS2 (–) +5 bp | intergenic (+63/-333) | 10% |
| **A1.4** | 44967 | *fixC* → | C→T | A263V (GCC→GTC) | 2% |
| | 345314 | *yahN* ← | +CA | coding (248/672 nt) | 2% |
| | 458014 | *clpX* → / → *lon* | IS*186* (+) +6 bp :: Δ1 bp | intergenic (+90/-93) | 7% |
| | 532261 | *allR* → | G→T | R9S (AGG→AGT) | 3% |
| | 847178 | *glnH* ← | G→A | A17V (GCG→GTG) | 2% |
| | 953901 | *focA* ← / ← *ycaO* | IS*5* (+) +4 bp | intergenic (-212/+191) | 2% |
| | 1195443 | *e14 prophage* | Δ15,192 bp | e14 prophage loss | 9% |
| | 1296579 | *adhE* ← | Δ3 bp | coding (764-766/2676 nt) | 1% |
| | 1720637 | *ydhK* → | Δ3 bp | coding (493-495/2013 nt) | 1% |
| | 1854063 | *ydjH* ← | G→T | T297N (ACC→AAC) | 5% |
| | 2076069 | *yeeW* → | G→A | pseudogene (106/168 nt) | 1% |
| | 2097981 | *gnd* ← | C→T | A438T (GCG→ACG) | 1% |
| | 2175243 | *gatY* ← / ← *fbaB* | T→G | intergenic (-17/+291) | 6% |
| | 2175295 | *gatY* ← / ← *fbaB* | IS*1* (–) +9 bp | intergenic (-69/+231) | 33% |
| | 2175297 | *gatY* ← / ← *fbaB* | IS2 (+) +5 bp | intergenic (-71/+233) | 91% |
| | 2827134 | *srlR* → | +GAAGA | coding (66/774 nt) | 3% |
| | 3216723 | *aer* ← | C→A | V126L (GTG→TTG) | 5% |
| | 3438154 | *rpoA* ← | G→A | L300F (CTT→TTT) | 26% |
| | 4540043 | *fimB* → / → *fimE* | IS*5* (–) +4 bp | intergenic (+461/-14) | 18% |
| | 4540060 | *fimE* → | IS2 (+) +5 bp | coding (1-5/597 nt) | 16% |
| | 4540184 | *fimE* → | IS*5* (–) +4 bp | coding (125-128/597 nt) | 6% |
| | 4540229 | *fimE* → | IS2 (+) +5 bp | coding (170-174/597 nt) | 10% |

| | | | | | |
|---|---|---|---|---|---|
| | 4540321 | *fimE* → | C→T | Q88* (CAG→TAG) | 2% |
| | 4540331 | *fimE* → | IS*5* (+) +4 bp | coding (272-275/597 nt) | 2% |
| | 4540389 | *fimE* → | IS*2* (+) +5 bp | coding (330-334/597 nt) | 10% |
| | 4540471 | *fimE* → | IS*5* (+) +4 bp | coding (412-415/597 nt) | 6% |
| | 4593394 | *yjjM* ← | A→G | F161L (TTT→CTT) | 4% |
| | 4638637 | *yjjY* → / → *yjtD* | IS*5* (+) +4 bp | intergenic (+72/-325) | 71% |
| **A1.5** | 4986 | *thrC* → | C→T | A418V (GCT→GTT) | 2% |
| | 458014 | *clpX* → / → *lon* | IS*186* (+) +6 bp :: Δ1 bp | intergenic (+90/-93) | 2% |
| | 720169 | *ybfK* → / ← *kdpE* | IS*1* (+) +8 bp | intergenic (+106/+103) | 2% |
| | 1195443 | *e14 prophage* | Δ15,192 bp | e14 prophage loss | 3% |
| | 1295061 | *adhE* ← | C→T | A762T (GCA→ACA) | 2% |
| | 1596359 | *yneO* ← / ← *lsrK* | Δ3 bp | intergenic (-348/+280) | 4% |
| | 2172381 | *gatB* ← | IS*5* (+) +4 bp | coding (205-208/285 nt) | 5% |
| | 2172834 | *gatA* ← | IS*1* (+) +9 bp | coding (230-238/453 nt) | 5% |
| | 2173262 | *gatZ* ← | IS*1* (+) +8 bp | coding (1075-1082/1263 nt) | 9% |
| | 2175267 | *gatY* ← / ← *fbaB* | G→T | intergenic (-41/+267) | 16% |
| | 2175295 | *gatY* ← / ← *fbaB* | IS*1* (–) +9 bp | intergenic (-69/+231) | 41% |
| | 2585982 | *acrD* → | G→A | V122V (GTG→GTA) | 2% |
| | 3265169 | *tdcA* ← / → *tdcR* | IS*5* (+) +4 bp | intergenic (-82/-230) | 9% |
| | 3532003 | *pck* → | Δ2 bp | coding (1164/1623 nt) | 2% |
| | 3653810 | *yhiD* ← | C→T | G39D (GGC→GAC) | 1% |
| | 3756149 | *selB* ← | C→T | G579D (GGC→GAC) | 4% |
| | 4166219 | *rrsB* → | C→A | noncoding (1538/1542 nt) | 10% |
| | 4635571 | *creC* → | Δ3 bp | coding (853-855/1425 nt) | 1% |
| | 4638628 | *yjjY* → / → *yjtD* | IS*2* (–) +5 bp | intergenic (+63/-333) | 64% |
| | 4638637 | *yjjY* → / → *yjtD* | IS*5* (+) +4 bp | intergenic (+72/-325) | 41% |
| | 4638670 | *yjjY* → / → *yjtD* | IS*5* (–) +4 bp | intergenic (+105/-292) | 8% |

| | | | | | |
|---|---|---|---|---|---|
| A2.1 | 627257 | *entB* → | C→T | P114L (CCA→CTA) | 3% |
| | 1140901 | *rne* ← | Δ3 bp | coding (2688-2690/3186 nt) | 2% |
| | 2175233 | *gatY* ← / ← *fbaB* | IS2 (+) +5 bp | intergenic (-7/+297) | 88% |
| | 2604462 | *hyfE* → | G→T | W60L (TGG→TTG) | 5% |
| | 2827091 | *srlR* → | Δ1 bp | coding (23/774 nt) | 7% |
| | 3734676 | *bax* ← | Δ3 bp | coding (523-525/825 nt) | 1% |
| | 4053901 | *glnL* ← | C→T | L154L (CTG→CTA) | 3% |
| | 4638670 | *yjjY* → / → *yjtD* | IS*5* (–) +4 bp | intergenic (+105/-292) | 1% |
| | 4638842 | *yjjY* → / → *yjtD* | +T | intergenic (+277/-123) | 94% |
| A2.3 | 118131 | *nadC* ← | Δ3 bp | coding (513-515/894 nt) | 1% |
| | 219229 | *tsaA* ← | G→T | G122G (GGC→GGA) | 2% |
| | 483856 | *acrA* ← | C→T | D330N (GAC→AAC) | 2% |
| | 1195443 | *e14 prophage* | Δ15,192 bp | e14 prophage loss | 7% |
| | 1325120 | *rluB* → | G→T | R82L (CGT→CTT) | 3% |
| | 1618295 | *eamA* ← | C→A | L289F (TTG→TTT) | 2% |
| | 1622428 | *ydeI* ← | G→T | P32T (CCG→ACG) | 2% |
| | 1720637 | *ydhK* → | Δ3 bp | coding (493-495/2013 nt) | 2% |
| | 1854063 | *ydjH* ← | G→T | T297N (ACC→AAC) | 4% |
| | 2079521 | *dacD* ← | T→A | I351F (ATT→TTT) | 3% |
| | 2172221 | *gatC* ← | IS*5* (+) +4 bp | coding (77-80/1356 nt) | 11% |
| | 2173905 | *gatZ* ← | Δ5 bp | coding (435-439/1263 nt) | 45% |
| | 2174867 | *gatY* ← | IS*4* (+) +12 bp | coding (349-360/855 nt) | 50% |
| | 2179265 | *yegV* → | G→A | V50I (GTT→ATT) | 3% |
| | 2403910 | *lrhA* ← | Δ5 bp | coding (750-754/939 nt) | 1% |
| | 2625564 | *yfgF* ← | G→T | A466D (GCC→GAC) | 3% |
| | 2808011 | *ygaZ* → | G→A | A125T (GCA→ACA) | 2% |
| | 2809759 | *emrA* → | C→T | A104V (GCC→GTC) | 2% |

| | | | | | |
|---|---|---|---|---|---|
| | 2827492 | *srlR* → | G→A | G142S (GGC→AGC) | 47% |
| | 3174366 | *cpdA* ← | Δ3 bp | coding (488-490/828 nt) | 3% |
| | 3249977 | *yqjG* → | T→C | I311T (ATT→ACT) | 1% |
| | 3681941 | *yhjK* ← | G→A | G567G (GGC→GGT) | 2% |
| | 3785915 | *envC* → | Δ3 bp | coding (1055-1057/1260 nt) | 1% |
| | 4176106 | *nusG* → | G→A | R114H (CGC→CAC) | 2% |
| | 4178685 | *rplL* → | Δ3 bp | coding (103-105/366 nt) | 1% |
| | 4540184 | *fimE* → | IS*5* (–) +4 bp | coding (125-128/597 nt) | 1% |
| | 4582042 | *hsdR* ← | C→A | D915Y (GAC→TAC) | 2% |
| | 4592986 | *yjjM* ← | IS*1* (–) +9 bp | coding (881-889/915 nt) | 1% |
| | 4601117 | *yjjP* ← / → *yjjQ* | IS*5* (+) +4 bp | intergenic (-236/-380) | 25% |
| | 4601140 | *yjjP* ← / → *yjjQ* | IS*5* (+) +4 bp | intergenic (-259/-357) | 13% |
| | 4601143 | *yjjP* ← / → *yjjQ* | IS*2* (–) +5 bp | intergenic (-262/-353) | 4% |
| | 4601148 | *yjjP* ← / → *yjjQ* | +G :: IS*5* (–) +3 bp | intergenic (-267/-350) | 2% |
| | 4638628 | *yjjY* → / → *yjtD* | IS*2* (–) +5 bp | intergenic (+63/-333) | 3% |
| | 4638637 | *yjjY* → / → *yjtD* | IS*5* (+) +4 bp | intergenic (+72/-325) | 5% |
| | 4638771 | *yjjY* → / → *yjtD* | IS*5* (+) +4 bp | intergenic (+206/-191) | 74% |
| **A2.4** | 210990 | *ldcC* → | G→T | E438* (GAG→TAG) | 3% |
| | 337004 | *yahF* → | C→T | R335C (CGT→TGT) | 3% |
| | 430331 | *yajD* → / ← *tsx* | A→T | intergenic (+155/+22) | 3% |
| | 953901 | *focA* ← / ← *ycaO* | IS*5* (+) +4 bp | intergenic (-212/+191) | 2% |
| | 1195443 | *e14 prophage* | Δ15,192 bp | e14 prophage loss | 4% |
| | 1708331 | *rsxG* → | C→T | A35V (GCT→GTT) | 2% |
| | 1720637 | *ydhK* → | Δ3 bp | coding (493-495/2013 nt) | 2% |
| | 1907380 | *kdgR* ← | +GC | coding (744/792 nt) | 1% |
| | 2172662 | *gatA* ← | IS*1* (+) +9 bp | coding (402-410/453 nt) | 86% |
| | 2172755 | *gatA* ← | IS*1* (–) +8 bp | coding (310-317/453 nt) | 5% |

| | | | | | |
|---|---|---|---|---|---|
| | 2172882 | *gatA* ← | IS*1* (+) +9 bp | coding (182-190/453 nt) | 2% |
| | 2173327 | *gatZ* ← | IS*1* (–) +8 bp | coding (1010-1017/1263 nt) | 4% |
| | 2636481 | *bamB* ← | G→A | P65L (CCG→CTG) | 2% |
| | 2827555 | *srlR* → | IS*3* (–) +5 bp :: Δ3 bp +T | coding (487-491/774 nt) | 4% |
| | 2827665 | *srlR* → | IS*150* (+) +3 bp | coding (597-599/774 nt) | 6% |
| | 3123436 | *glcF* ← | Δ2 bp | coding (46/1224 nt) | 3% |
| | 3734676 | *bax* ← | Δ3 bp | coding (523-525/825 nt) | 1% |
| | 4178685 | *rplL* → | Δ3 bp | coding (103-105/366 nt) | 1% |
| | 4364872 | *dcuA* ← / ← *aspA* | IS*4* (+) +12 bp | intergenic (-76/+31) | 3% |
| | 4442468 | *tamB* → | Δ1 bp | coding (334/3780 nt) | 2% |
| | 4540184 | *fimE* → | IS*5* (+) +4 bp | coding (125-128/597 nt) | 15% |
| | 4540331 | *fimE* → | IS*5* (+) +4 bp | coding (272-275/597 nt) | 48% |
| | 4638628 | *yjjY* → / → *yjtD* | IS*2* (–) +5 bp | intergenic (+63/-333) | 78% |
| | 4638771 | *yjjY* → / → *yjtD* | IS*5* (+) +4 bp | intergenic (+206/-191) | 2% |
| **A2.5** | 223673 | *gmhB* → / → *rrsH* | A→G | intergenic (+265/-98) | 20% |
| | 923583 | *clpA* → | Δ1 bp | coding (1097/2277 nt) | 3% |
| | 953892 | *focA* ← / ← *ycaO* | IS*5* (+) +13 bp | intergenic (-203/+191) | 22% |
| | 953896 | *focA* ← / ← *ycaO* | IS*5* (+) Δ5 bp | intergenic (-207/+195) | 19% |
| | 1195443 | *e14 prophage* | Δ15,192 bp | e14 prophage loss | 2% |
| | 1198930 | *intE* ← | C→G | G367A (GGT→GCT) | 47% |
| | 1292419 | *hns* ← / → *tdk* | IS*5* (+) +4 bp | intergenic (-274/-328) | 8% |
| | 1299320 | *oppA* → | C→T | Q39* (CAA→TAA) | 22% |
| | 1720637 | *ydhK* → | Δ3 bp | coding (493-495/2013 nt) | 2% |
| | 2172755 | *gatA* ← | IS*1* (–) +8 bp | coding (310-317/453 nt) | 56% |
| | 2816091 | *argZ* ← | G→T | noncoding (67/77 nt) | 3% |
| | 2827297 | *srlR* → | A→T | K77* (AAG→TAG) | 26% |
| | 2827492 | *srlR* → | G→A | G142S (GGC→AGC) | 49% |

| | | | | | |
|---|---|---|---|---|---|
| | 3123436 | *glcF* ← | Δ2 bp | coding (46/1224 nt) | 4% |
| | 3253058 | *yhaK* → / → *yhaL* | Δ3 bp | intergenic (+16/-5) | 2% |
| | 3453566 | *gspA* ← / → *gsp C* | +ACCC | intergenic (-146/-34) | 34% |
| | 3881169 | *dnaA* ← | Δ1 bp | coding (584/1404 nt) | 2% |
| | 4048535 | *yihA* ← | G→A | A85V (GCG→GTG) | 2% |
| | 4053440 | *glnL* ← | C→T | R308H (CGC→CAC) | 3% |
| | 4072821 | *yihW* → | Δ74 bp | coding (130-203/786 nt) | 1% |
| | 4178685 | *rplL* → | Δ3 bp | coding (103-105/366 nt) | 1% |
| | 4601134 | *yjjP* ← / → *yjjQ* | IS*5* (–) +4 bp | intergenic (-253/-363) | 4% |
| | 4601187 | *yjjP* ← / → *yjjQ* | IS*1* (–) +8 bp | intergenic (-306/-306) | 2% |

**Table S4. Functional Category of Mutational Targets With a Frequency Above 5% Observed in *E. coli* Populations After 24 Days of Evolution in Very Old Mice. Related to Figures 2 and 3.** Summary of the mutational targets above 5% identified in *E. coli* populations after 24 days of evolution in the gut of very old mice. The table includes the following information for each mutational target: gene function (from STRING [S6] and EcoCyc [S7]), corresponding KEGG pathways, Gene Ontology (GO) Biological Processes, and the functional category assigned (metabolism, stress response, or others). The table also highlights overlap in mutational targets between different age groups.

| | | | | **Mutated in** | | |
|---|---|---|---|---|---|---|
| **Mutational target** | **Function** | **KEGG pathway / GO Biological Process** | **Attributed category** | **Young** | **Old** | **Very Old** |
| *gat* operon | Galactitol metabolic process | Galactose metabolism / Galactitol metabolic process | Metabolism | √ | √ | √ |
| *srlR* | Negative regulation of DNA-templated transcription - represssion of an operon involved in the transport and utilization of sorbitol | Fructose and mannose metabolism / Sorbitol transmembrane transport | Metabolism | √ | √ | √ |
| *tdcA / tdcR* | Transport and metabolism of threonine and serine | und. / L-threonine catabolic process to propionate | Metabolism | √ | √ | √ |

| | | | | | | |
|---|---|---|---|---|---|---|
| *hns* | Regulation of several genes, including in response to environmental changes and adaptation to stress | und. / Bacterial nucleoid packaging | Stress | √ | √ | √ |
| *ydiF* | Acetate CoA-transferase (predicted based on sequence similarity) | Aminobenzoate/benzoate/lysine degradation / Fatty acid metabolism | Metabolism | √ | √ | |
| *yjjY / yjtD* | ***arcA*** (gene affected by these mutations): response to changing respiratory conditions of growth with a prominent role in the anaerobic repression of genes associated with aerobic metabolism | Two-component system / Protein autophosphorylation | Stress | | √ | √ |
| *lrhA* | Global transcriptional regulator, including of genes involved in biofilm formation, motility, chemotaxis, flagellum synthesis, and stress response | und. / Positive regulation of transcription, DNA-templated | Stress | | √ | √ |
| *iscR ** | Iron-sulfur cluster negative regulator | und. / Protein maturation by [4Fe-4S] cluster transfer | Stress | | √ | √ |
| *gfcE ** | Putative O-antigen capsule outer membrane auxillary protein export channel; May be involved in polysaccharide transport | Two-component system / Peptidyl-tyrosine autophosphorylation | Others | | √ | √ |
| *sel* operon *(selA,selB)* | ***selA***: selenocysteine synthase; ***selB***: selenocysteinyl-tRNA-specific translation factor necessary for the incorporation of selenocysteine into proteins | und. / Selenocysteine biosynthetic process | Metabolism | | √ | √ |
| *fim* operon *(fimE / fimA or fimE)* | Both genes from the *fim* operon, responsible for the synthesis of type 1 fimbriae | und. / Cell adhesion involved in single-species biofilm formation | Stress | | √ | √ |
| *yjjP / yjjQ* | ***yjjP***: implicated in succinate transport; ***yjjQ***: transcriptional repressor of genes required for flagellar synthesis, capsule formation, and other genes related to virulence | ***yjjP***: und. / Succinate transport; ***yjjQ***: und. / und. | Metabolism / Stress | √ | | √ |
| *focA / ycaO* | ***focA***: bidirectional formate transporter, regulates intracellular formate levels during anaerobic respiration and mixed acid fermentation; ***ycaO***: involved in the β-methylthiolation of 30S ribosomal subunit protein S12, molecular function unknown | ***focA***: Butanoate/pyruvate metabolism / und.; ***ycaO***: und. / Macromolecule modification | Metabolism / Metabolism | √ | | √ |

| | | | | | | |
|---|---|---|---|---|---|---|
| *kdgR* | Repression of transcription of the operons involved in transport and catabolism of 2-keto-3-deoxy gluconate | und. / D-glucuronate catabolic process | Metabolism | √ | | √ |
| *yjjM* | Predicted transcriptional regulator that is essential for growth on L-galactonate as the sole carbon source | und. / und. | Metabolism | √ | | √ |
| *yfjL / yfjM ** | ***yfjL***: impairment of phage DNA synthesis ***yfjM***: uncharacterized protein | ***yfjL***: und. / Defense response to virus; ***yfjM***: und. / Defense response to virus | Stress / Stress | √ | | √ |
| *dcuB / dcuR* | ***dcuB***: fumarate/succinate antiporter during anaerobic fumarate respiration; ***dcuR***: Involved in the C4-dicarboxylate-stimulated regulation of the genes encoding the anaerobic fumarate respiratory system | ***dcuB***: Two-component system / Fermentation; ***dcuR***: Two-component system / Phosphorelay signal transduction system | Metabolism / Metabolism | √ | | |
| *yeaR* | Transcription is induced in response to nitrate, nitrite, and nitric oxide; involved in xenobiotic metabolic processes | und. / und. | Stress | √ | | |
| *eutH / eutG* | ***eutH:*** possible ethanolamine transport; ***eutG***: ethanol dehydrogenase involved in ethanolamine utilization | ***eutH***: Glycerophospholipid metabolism / Ethanolamine catabolic process; ***eutG***: Naphthalene degradation / Ethanol metabolic process | Metabolism / Metabolism | √ | | |
| *garK* | Catalyzation the formation of 2-phosphoglycerate from D-glycerate | Glycine, serine and threonine metabolism / Organic acid phosphorylation | Metabolism | √ | | |
| *atoC* | Stimulation of the expression of the atoDAEB operon, leading to short chain fatty acid catabolism and activation of the poly-(R)-3-hydroxybutyrate (cPHB) biosynthetic pathway; Also involved in the regulation of motility and chemotaxis, via transcriptional induction of the flagellar regulon | Aminobenzoate degradation / Bacterial-type flagellum-dependent swimming motility | Metabolism | √ | | |

| | | | | |
|---|---|---|---|---|
| *codA / cynR* | ***codA***: Catalyzation of the deamination of cytosine to uracil, which allows the cell to utilize cytosine for pyrimidine nucleotide synthesis; ***cynR***: Transcriptional activator of cyn operon, involved in cyanate metabolism | ***codA:*** Pyrimidine metabolism / Cytosine metabolic process; ***cynR:*** und. / Cyanate catabolic process | Metabolism / Metabolism | √ |
| *ybcN* | Encodes the cryptic lambdoid prophage DLP12 and is a distantly related member of the Orf phage protein family | und. / und. | Others | √ |
| *sppA* | Digests cleaved signal peptides *in vitro; in vivo* function is currently unknown | und. / Proteolysis | Others | √ |
| *yobB* | Poorly characterized; C-N hydrolase family protein | und. / und. | Others | √ |
| *nrdA / nrdB* | ***nrdA*** and ***nrdB****:* Providing of the precursors necessary for DNA synthesis | ***nrdA*** and ***nrdB***: Pyrimidine metabolism / Nucleoside diphosphate biosynthetic process | Metabolism / Metabolism | √ |
| *syd* | Possibly serves as a quality control system for assembly of the SecY secretion channel | und. / und. | Others | √ |
| *sodA / kdgT* | ***sodA***: superoxide dismutase, implicated in the response to a large number of environmental changes that lead to the generation of superoxide radicals; ***kdgT***: 2-keto-3-deoxy-D-gluconate transporter | ***sodA***: Tryptophan metabolism / Hydrogen peroxide catabolic process; ***kdgT***: und. / D-glucuronate catabolic process | Stress / Metabolism | √ |
| *aidB* | DNA alkylation damage repair protein; Part of the adaptive DNA-repair response to alkylating agents | Caprolactam degradation / DNA dealkylation involved in DNA repair | Stress | √ |
| *arcA* | Response to changing respiratory conditions of growth with a prominent role in the anaerobic repression of genes associated with aerobic metabolism | Two-component system / Protein autophosphorylation | Stress | √ |

| | | | | | |
|---|---|---|---|---|---|
| *mltB_srlA* | ***mltB***: Membrane-bound lytic murein transglycosylase B; Murein-degrading enzyme. Catalyzes the cleavage of the glycosidic bonds between N-acetylmuramic acid and N-acetylglucosamine residues in peptidoglycan; ***srlA***: Glucitol/sorbitol-specific enzyme IIC component of PTS, a major carbohydrate active transport system | ***mltB:*** und. / Cell wall macromolecule catabolic process; ***srlA***: Phosphotransferase system (PTS) / Sorbitol transmembrane transport | Metabolism / Metabolism | | √ |
| *thrB* | Homoserine kinase; Catalyzation of the ATP-dependent phosphorylation of L-homoserine to L-homoserine phosphate | Monobactam biosynthesis / Homoserine biosynthetic process | Metabolism | | √ |
| *ykfK* | Poorly characterized; CP4-6 prophage | und. / und. | Others | | √ |
| *ddlA_iraP* | ***ddlA***: D-alanine-D-alanine ligase A; Cell wall formation. Potentially involved in resistance to X-ray radiation and vancomycin; ***iraP***: one of several small anti-adaptor proteins; it is required for stabilization of the alternative sigma factor σS during phosphate starvation and plays a role in stationary phase and nitrogen starvation | ***ddlA***: D-Alanine metabolism, Vancomycin resistance / D-alanine biosynthetic process;***iraP***: und. / Negative regulation of protein catabolic process, Cellular response to starvation | Stress / Stress | | √ |
| *mngB_cydA* | ***mngB***: Poorly characterized alpha-mannosidase; May hydrolyze 6-phospho-mannosyl-D-glycerate to mannose-6- phosphate and glycerate;***cydA***: Cytochrome d terminal oxidase, subunit I; It is the component of the aerobic respiratory chain of that predominates when cells are grown at low aeration | ***mngB***: und. / und.;***cydA***: Oxidative phosphorylation / Aerobic electron transport chain | Others / Stress | | √ |
| *puuE* | A putrescine-inducible GABA aminotransferase, which allows cells to grow on putrescine as the sole source of nitrogen. Its activity is induced by growth on putrescine and repressed by addition of succinate and under conditions of limited oxygen availability | Beta-alanine metabolism / Glutamate catabolic process | Stress | | √ |
| *spoT* | A key enzyme that acts as a mediator of the stringent response which coordinates a variety of cellular activities in response to changes in nutritional abundance | Purine metabolism / Guanosine tetraphosphate biosynthetic process | Stress | | √ |

| | | | | | | |
|---|---|---|---|---|---|---|
| *clpX_lon* | ***clpX***: an ATP-dependent molecular chaperone that serves as a substrate-specifying adaptor for the ClpP serine protease in the ClpXP and ClpAXP protease complexes; ***lon***: an ATP-dependent protease responsible for degradation of misfolded proteins and several rapidly degraded regulatory proteins. Required for cellular homeostasis and for survival from DNA damage and stress-induced developmental changes | ***clpX***: und. / Mitochondrial protein processing***;*** ***lon***: und. / Protein denaturation | Metabolism / Stress | | | √ |
| *allR* | Allantoin repressor; regulates several genes involved in the anaerobic utilization of allantoin as a nitrogen source | und. / Allantoin catabolic process | Metabolism | | | √ |
| *gmhB_rrsH* | ***gmhB***: D,D-heptose 1,7-bisphosphate phosphatase of the ADP-heptose biosynthesis pathway; ***rrsH***: Ribosomal operon of the 16S rRNA (the RNA component of the small subunit (30S subunit) of the *E. coli* ribosome) | ***gmhB:*** und. / und.; r***rsH***: Ribosome / und. | Metabolism / Others | | | √ |
| *oppA* | Periplasmic binding protein of an ABC transporter that mediates the high affinity uptake of oligopeptides; Predicted to play a role in the heat shock response | Beta-Lactam resistance / Oligopeptide import across plasma membrane | Stress | | | √ |
| *fdoG* | α subunit of formate dehydrogenase O; Allows the use of formate as major electron donor during anerobic respiration | Methane metabolism / Heme oxidation | Metabolism | | | √ |
| *intE* | Integrase from the cryptic lambdoic prophage e14, necessary for integration and excision of the phage into the host genome | und. / Establishment of integrated proviral latency | Others | | | √ |
| *hns_tdk* | ***hns***: Regulation of several genes, including in response to environmental changes and adaptation to stress; ***tdk***: Deoxythymidine kinase, it catalyzes the ATP-dependent conversion of deoxythymidine to dTMP; it is an enzyme of the salvage pathways of pyrimidine deoxyribonucleotides | ***hns***: und. / Bacterial nucleoid packaging; ***tdk***: Pyrimidine metabolism / Pyrimidine deoxyribonucleoside metabolic process | Stress / Metabolism | | | √ |

| | | | | | | |
|---|---|---|---|---|---|---|
| *rpoA* | The α subunit of the RNA polymerase core enzyme. This subunit plays an important role in subunit assembly since its dimerization is the first step in the sequential assembly of subunits to form the holoenzyme | Ribosome / DNA-templated transcription | Others | | | √ |
| *gspA / gspC* | ***gspA*** and ***gspC***: Part of a cluster of genes (gsp) predicted to encode the components of a type II secretion system for the export of proteins. | Bacterial secretion system / Protein secretion by the type II secretion system | Others | | | √ |
| *rrsB* | Ribosomal operon of the 16S rRNA (the RNA component of the small subunit (30S subunit) of the *E. coli* ribosome) | Ribosome / und. | Others | | | √ |
| *nhaA* | An Na+:H+ antiporter with a prominent role in sodium ion and alkaline pH homeostasis; supports the maintenance of cytoplasmic pH under alkali challenge; also has an important role in maintaining antibiotic tolerance under starvation, and is upregulated during nutrient starvation | und. / Response to alkaline pH | Stress | | | √ |
